# Small heat shock protein HSPB8 interacts with a pre-fibrillar TDP43 low complexity domain species to delay fibril formation

**DOI:** 10.1101/2025.01.28.635368

**Authors:** Khaled M. Jami, Katherine Corbett, Daniel C. Farb, Kayla M. Osumi, Catelynn C. Shafer, Sophie Criscione, Dylan T. Murray

**Affiliations:** Department of Chemistry, University of California, One Shields Avenue, Davis, California, 95616, United States of America; Department of Molecular and Cell Biology, University of Connecticut, 91 North Eagleville Road, Storrs, Connecticut, 06269, United States of America

**Author notes:** Correspondence: Dylan T. Murray.

**Keywords:** protein aggregation, liquid droplet, small heat shock protein, protein fibril, RNA-binding protein, biomolecular condensate, ATP-independent chaperone

## Abstract

The loss of cellular proteostasis through aberrant stress granule formation is implicated in neurodegenerative diseases. Stress granules are formed by biomolecular condensation involving protein-protein and protein-RNA interactions. These assemblies are protective, but can rigidify, leading to amyloid-like fibril formation, a hallmark of the disease pathology. Key proteins dictating stress granule formation and disassembly, such as TDP43, contain low-complexity (LC) domains that drive fibril formation. HSPB8, a small heat shock protein, localizes to stress granules, has known aggregation delaying activity, and helps direct aggregated proteins to protein degradation pathways. It is not known how HSPB8 interacts with aggregation prone LC domains in stress granules. Here, we examined the interaction between isolated HSPB8 and the TDP43 LC using thioflavin T (ThT) and fluorescence polarization (FP) aggregation assays, fluorescence microscopy and photobleaching experiments, and crosslinking mass spectrometry (XL-MS). Our results indicate that HSPB8 delays TDP43 LC aggregation through domain-specific interactions with fibril nucleating species, without affecting fibril elongation rates. These findings provide mechanistic insight into how HSPB8 mediates LC domain aggregation and provides bases for investigating how the TDP43 LC subverts chaperone activity in neurodegenerative disease and differing mechanisms between members of the HSPB protein family.

## Introduction

A mechanistic understanding of how cells lose the ability to regulate stress granule formation could provide new treatment modalities for neurodegenerative diseases such as amyotrophic lateral sclerosis (ALS), frontotemporal dementia (FTD), and Alzheimer’s Disease (AD). Stress granules are cytoplasmic membraneless organelles formed by transient multivalent interactions between proteins, and between proteins and RNA (1). These organelles are thought to be cytoprotective, arresting translational machinery in response to various cellular stresses such as heat shock (2), viral infection (3), and oxidative stress (4).

Many of the proteins driving stress granule and other biomolecular condensation processes have similar functional and structural motifs, often containing RNA-binding domains and an intrinsically disordered low-complexity (LC) domain (5, 6) biased toward a subset of the 20 naturally occurring amino acids. These LC regions drive biomolecular condensation through transient protein-protein interactions (PPI), especially self-association (7), forming stress granules with fluid liquid droplet properties (8, 9). The formation of the stress granules is reversible, however under prolonged stress, the constituent proteins can lose fluidity and form more rigid assemblies. The rigidification process necessarily sequesters and concentrates aggregation-prone LC proteins and increases the likelihood of forming amyloid-like fibrils or other proteinaceous aggregates (10, 11). Indeed, this molecular aging process is a putative mechanism by which the aggregated foci observed in neurodegenerative disease form (11).

Amyloid-like fibrils and aggregates composed of full-length TDP43 and C-terminal truncations including the LC domain (6, 12, 13) have been found in an increasing number of neurodegenerative diseases, including ALS (14), FTD (15), and the recently classified limbic-predominant age-related TDP43 encephalopathy (LATE) (16). TDP43 is an RNA and DNA binding protein with activities in translation, transcription, and splicing (17). Along with many RNA-binding proteins (RBPs), TDP43 is a major constituent and regulator of stress granules (18–20). Previous work has demonstrated that in vitro TDP43 liquid droplets have properties akin to cellular stress granules (21–23). In addition, aged liquid droplets consisting of TDP43 LC (24) and TIA1 LC (25) domains form amyloid-like fibrils, further emphasizing the possible link between stress granule aging and pathogenesis.

Crucially, evidence is growing that stress granules are regulated by ATP-dependent molecular chaperones like HSP70 (26), HSP90 (27), and their counterparts the ATP-independent, small heat shock proteins (sHSP). sHSP can localize inside stress granules (28, 29) and stabilize aggregation-prone proteins within stress granules (30). Furthermore, sHSP harbor mutations linked to inherited neuropathies, such as Charcot-Marie-Tooth (CMT) disease and distal hereditary motor neuropathy (dHMN) (31, 32).

HSPB8 is an sHSP that engages in various activities during detrimental proteosome inhibition (33). It is an important component of the protein quality control system where it prevents irreversible aggregation and assists in the degradation of misfolded proteins through recruitment of co-chaperones (26, 34). Additionally, in vivo studies on primary motor neurons (35) and a *Drosophila melanogaste*r model of ALS (36) show that high HSPB8 expression levels are associated with the absence of aggregated TDP43 and reduces TDP43 induced toxicity.

Despite their importance, a complete mechanistic description of how sHSP interact with aggregation-prone LC domains in the context of stress granule fluidity and persistence, and amyloid-like fibril formation does not exist. We focus on the C-terminal LC domain of TDP43 (residues 262–414; Fig. 1, top), since neurodegenerative disease related aggregation involves C-terminal truncations of TDP43 (37), the segment that forms pathogenic fibrils in patient tissues resides in the LC domain (38, 39), and experimental and computational studies from other laboratories have focused on the TDP43 LC domain (40–44). In this work, we dissect the PPI between the TDP43 LC domain and HSPB8 through thioflavin T (ThT) and fluorescence polarization (FP) aggregation assays, fluorescence microscopy and photobleaching experiments, and chemical crosslinking paired with mass spectrometry (XL-MS). ThT and FP assays report on aggregation and binding kinetics and points to a possible mechanism of inhibition. Fluorescence microscopy experiments monitor changes in TDP43 LC liquid droplet fluidity in the presence of HSPB8. Finally, XL-MS details a putative site of binding between HSPB8 and TDP43 LC. These findings offer insights into the mechanism by which sHSP mitigate protein aggregation, directing future efforts to determine how aggregation-prone RNA-binding proteins like TDP43 evade chaperone activity in neurodegenerative diseases.

**Figure 1.**
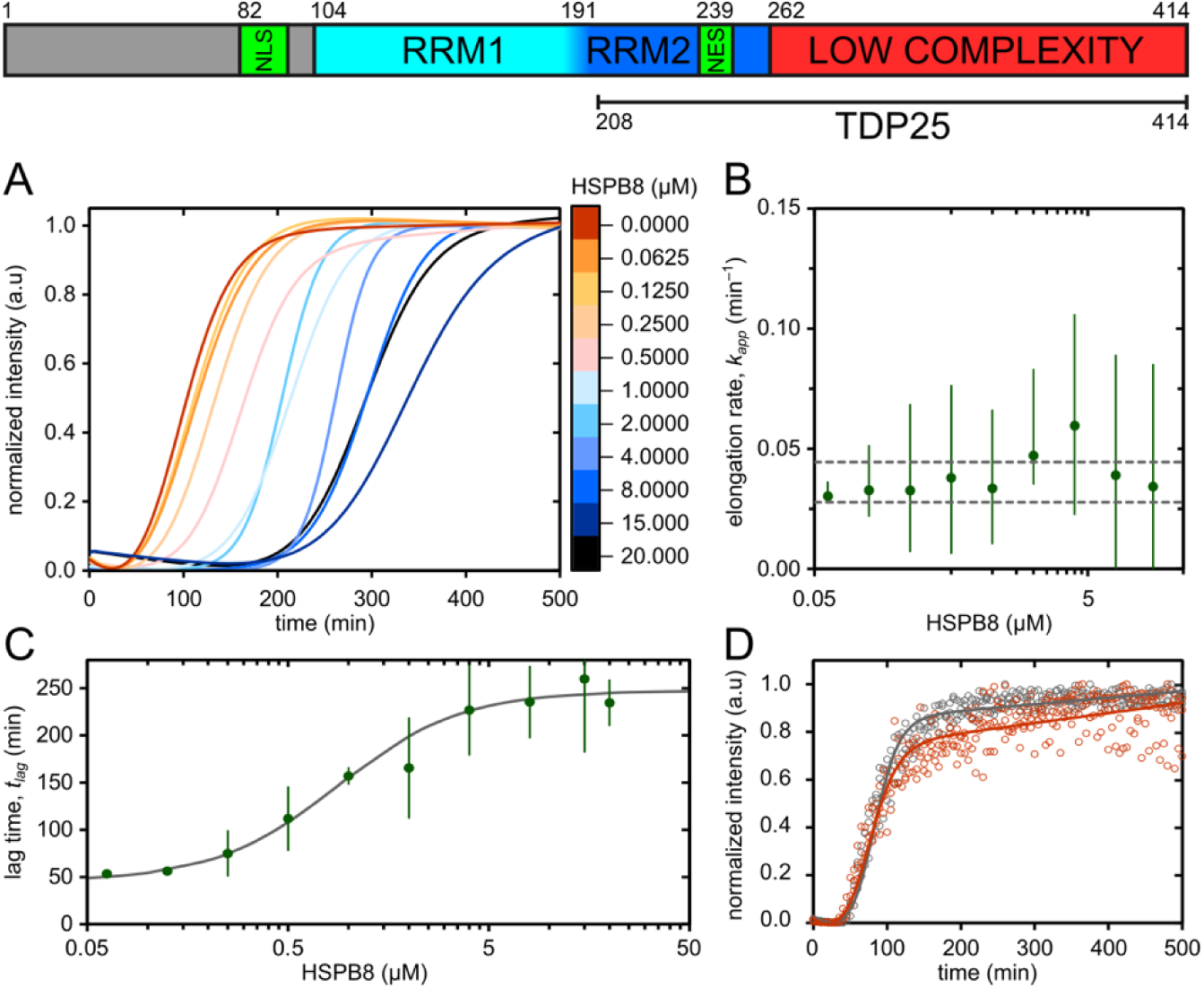
HSPB8 delays aggregation of TDP43 LC by preventing nucleation. Top, TDP43 domain architecture. Nuclear localization sequence (NLS), RNA recognition motifs (RRM), and nuclear export sequence (NES) domains are indicated with residues numbers at the top of the cartoon. The LC domain used in this study is residues 262–414. The C-terminal fragments (TDP25) found in neurodegenerative disease derive from enzymatic cleavage at residue 208. (A) Normalized, best-fit curves to ThT fluorescence data from 20 μM TDP43 LC aggregation assays with varying HSPB8 concentrations. Curves are generated using the average of the best-fit lag time (*t_lag_*) and apparent elongation rate (*k_app_*) parameters from n = 3 measurements. (B) A plot of the average *k_app_* as a function of HSPB8 concentration, p = 0.467 (Kruskal-Wallis test, α = 0.05). Error bars represent the 95% confidence interval. Horizontal dashed gray lines represent the 95% confidence interval for the TDP43 LC only condition (0 μM HSPB8) shown in red in panel (A). (C) A plot of *t_lag_*as a function of HSPB8 concentration. A best-fit sigmoidal curve calculated using a modified Hill equation is shown in gray. Error bars correspond to ±1 standard deviation. (D) Normalized ThT fluorescence data and best-fit curves for TDP43 LC aggregation assays with 8 µM BSA (black) and without (red). Curves are generated using the average of the best-fit *t_lag_*and *k_app_* parameters from n = 3 measurements. All experimental measurements are shown as circles in their respective color.

## Results

### HSPB8 Delays TDP43 LC Fibril Nucleation

Fig. 1 shows TDP43 domain architecture. For the experiments described here, residues 262–414 (TDP43 LC) were recombinantly expressed in E. coli with a His-tag and purified using metal affinity chromatography, enzymatic cleavage of the His-tag, and cation exchange chromatography. To investigate how the chaperone, HSPB8, modulates the assembly kinetics of the TDP43 LC, we first employed a ThT-fluorescence aggregation assay. ThT assays are ideal for detecting amyloid-like protein fibrils and studying their formation kinetics. The use of ThT to monitor TDP43 LC aggregation behavior is well-characterized (24, 45), providing a robust baseline for kinetic perturbation studies.

Fig. 1A and Supplemental Fig. S1 show a pronounced HSPB8 concentration-dependent change in the aggregation behavior of TDP43 LC measured by ThT fluorescence. TDP43 LC aggregation curves were modeled using the approach described by Malmos et al. (46) to extract the apparent rate constant of fibril elongation (*k_app_*) and the lag time required to accumulate a critical concentration of fibril-forming nuclei (*t_lag_*). Fig. 1B shows that under the physiological ionic strength and pH conditions used in the aggregation assay, increasing HSPB8 concentrations did not significantly affect *k_app_*. However, Fig. 1C shows the dependence of *t_lag_* on HSPB8 concentration is consistent with a hyperbolic curve, expected for saturation binding phenomena. Nonlinear regression analysis using a modified Hill Equation provided a measure of the ability of HSPB8 to delay the onset of TDP43 LC aggregation, with a half maximal inhibitory concentration (IC_50_) of 0.84 ± 0.36 µM. Note here the maximal inhibition is the maximum *t_lag_* time for a given TDP43 LC concentration at a set ionic strength and pH. This IC_50_ value indicates moderate affinity between HSPB8 and a species of TDP43 LC present during the aggregation process. This value aligns with previously measured dissociation constants for other sHSP, which are on the order of a few µM for HSPB5 interacting with alpha-synuclein and the beta-amyloid peptide (47–49). The Hill coefficient of 1.4 ± 0.2 suggests no or minimal cooperativity within experimental uncertainty.

HSPB8-dependent lengthening of *t_lag_* with no effect on *k_app_* suggests that HSPB8 delays TDP43 LC aggregation by interacting with fibril competent nuclei or unfolded soluble proteins, rather than with fibrils or another species contributing to ThT fluorescence. Once nucleation occurs, fibril propagation continues unaffected by HSPB8. Fig. 1D shows that non-specific PPI or crowding were not the cause of the observed effect, as the addition of bovine serum albumin (BSA) as a decoy protein produced no impact on the ThT profile for the TDP43 LC under aggregating conditions.

### HSPB8 Interacts with Prefibrillar TDP43 LC Species

We hypothesized that the HSPB8 substrate might be a species appearing early in the TDP43 aggregation process, such as soluble or pre-aggregated, nucleation-prone protein. Binding was qualitatively assessed by matrix-assisted laser desorption/ionization mass spectrometry with a time-of-fight mass analyzer (MALDI-TOF). Supplemental Fig. S2 shows several low intensity peaks unique to the HSPB8-TDP43 LC mixture consistent with 1^+^, 2^+^, and 3^+^ charged states for a 1:1 complex of HSPB8 and TDP43 LC, suggesting a possible weak PPI between soluble TDP43 LC and HSPB8.

To quantify and directly assess the interaction between HSPB8 and TDP43 LC, we then measured the fluorescence polarization (FP) of fixed amounts of fluorescently labeled HSPB8 with a range of soluble TDP43 LC concentrations. Fig. 2A shows that immediately after mixing (i.e. t = 0), there was a minimal change in FP with increasing concentrations of TDP43 LC. This observation is inconsistent with the MALDI-TOF measurements showing a complex of soluble TDP43 LC and HSPB8. However, the net charge on HSPB8 and TDP43 LC are predicted to be opposite based on calculated isoelectric points (HSPB8, pI = 5.5; TDP43 LC, pI = 10.6). Thus, gas-phase ion-pairing of HSPB8 and TDP43 LC is possible under our pH 7.2 MALDI-TOF measurement conditions. The observation is also inconsistent with HSPB8 binding to natively unfolded TDP43 LC in solution, as a concentration dependent effect would be observed at t = 0 for such a case. The initial FP measurements therefore suggest that the primary substrate of HSPB8 must form as TDP43 LC aggregation progresses.

**Figure 2.**
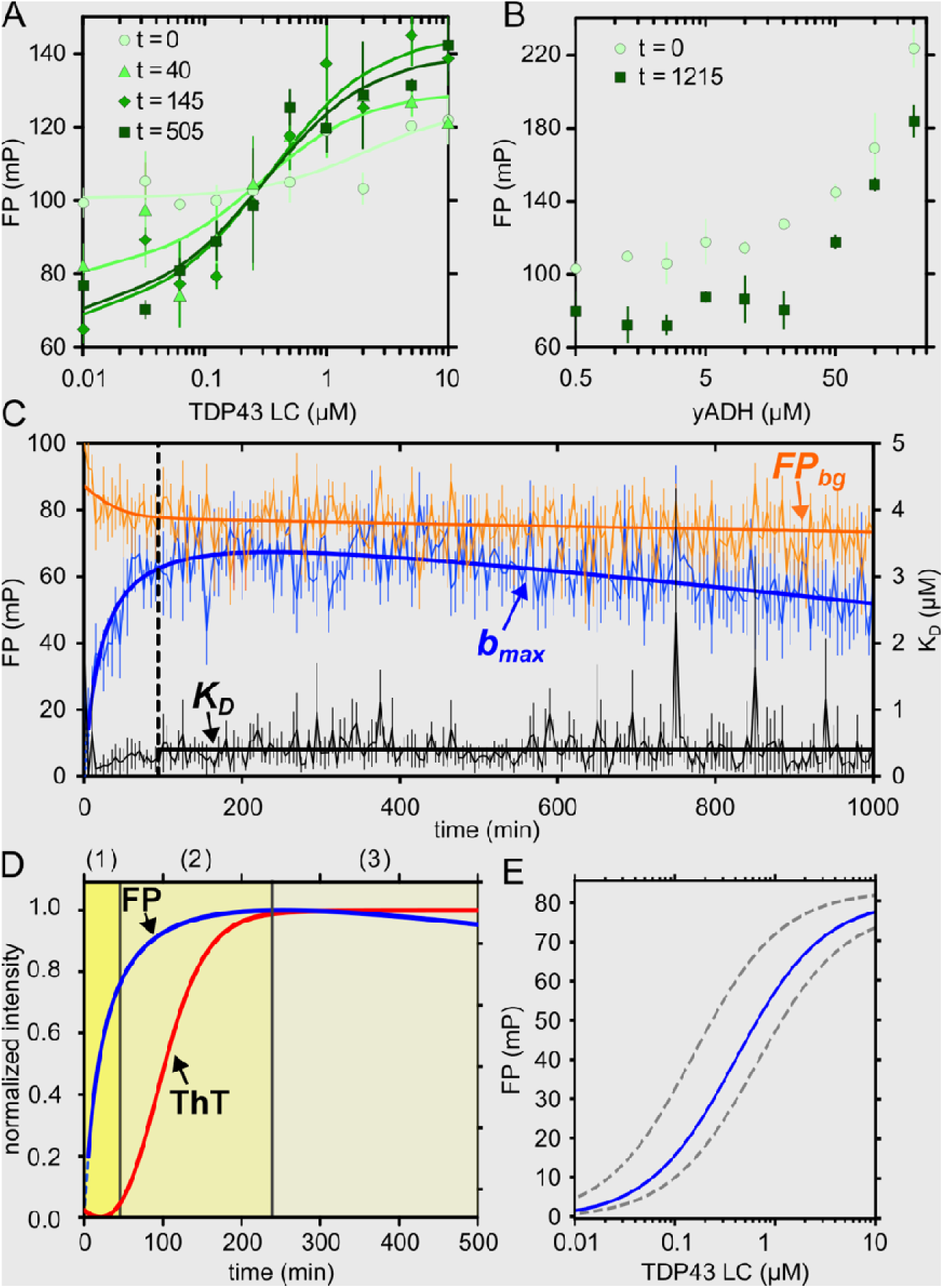
Fluorescence polarization monitors HSPB8 chaperone activity with pre-fibrillar TDP43 LC. (A) Fluorescence polarization (FP) curves at various time points during a TDP43 LC aggregation assa with 5 μM fluorescein-labeled HSPB8. Curves are plotted using the average best-fit parameters from n = 3 measurements. (B) FP values for the yADH aggregation assay before (t = 0 min) and after temperature induced aggregation (t = 1215 min) with 5 μM fluorescein-labeled HSPB8. For (A) and (B), all points are the average of n = 3 measurements, error bars represent ±1 standard deviation, and legend values are all in minutes. (C) Plot of the time dependence of the fitting parameters from the FP aggregation assay. The thin jagged-lines represent the best-fit parameter values obtained at each time point for the apparent background fluorescence (*FP_bg_*, orange), the apparent maximum number of TDP43 LC species capable of interacting with HSPB8 (*b_max_*, blue), and the apparent dissociation constant (*K′_D_*, black). The error bars correspond to ± 1 standard deviation from n = 3 measurements. The vertical black dashed line is at 94 min where *K_D_* averaging begins. The thick smooth lines are the optimized functions describing how the binding model parameters evolve over time. (D) Plot of the best-fit curves for the normalized FP *b_max_* (blue) and normalized ThT (red) data. The ThT assay was recorded in parallel to the FP assay using 10 µM TDP43 LC, 5 µM HSPB8, and 20 µM ThT. The three highlighted areas correspond to the (1) lag or nucleation phase ending at *t_lag_* (45 min), (2) elongation or growth phase ending at the maxima of the *b_max_* curve (238 min), and (3) plateau phase. (E) A background corrected FP plot generated using the fitted curves from all three separate plots in (C). The blue and orange dashed lines represent the variation in FP dictated by the standard deviation from the least-squares fit for *K_D_* and *B_max_*.

To probe transient processes in the TDP43 LC aggregation process, such as the formation of fibril-forming nuclei, the FP-based aggregation assay using fluorescein-labeled HSPB8 was monitored over time. Extensive characterization during our optimization of TDP43 LC sample preparation indicates that with agitation, the TDP43 LC aggregation parameters (*t_lag_* and *k_app_*) are concentration-independent. Therefore, FP results can be compared across varying TDP43 LC concentrations. While Fig. 2A shows little change in HSPB8 FP as a function of TDP43 LC concentration at initial time points, after 40 min the FP response is consistent with binding described by a non-cooperative Hill equation.

Fig. 2B shows a similar experiment with yeast alcohol dehydrogenase (yADH). The FP values did not plateau within the experimental concentration range, indicating that the binding of HSPB8 to yADH aggregating species is much weaker. Although yADH aggregation produced a larger change in overall HSPB8 FP, the significantly larger mass of yADH, existing as a tetramer of roughly 150 kDa (50), compared to 15.5 kDa TDP43 LC monomers, dictates the polarization response should be larger for HSPB8 binding to yADH than TDP43 LC. These results are intriguing given yADH’s common use as a benchmark protein for studying chaperone activity, including HSPB8 (51–53). Nevertheless, Supplemental Fig. S3A shows our preparations of HSPB8 exhibit the expected chaperone activity on yADH aggregation triggered by a reducing agent.

Fig. 2C shows the best-fit parameters for the HSPB8 FP response as a function of TDP43 LC concentration varied during the ∼16 h experiment. These parameters are *b_max_*, a measure of the number TDP43 LC species capable of binding to HSPB8; *FP_bg_*, the background fluorescence signal; and *K_D_*, the dissociation constant for TDP43 LC binding to HSPB8. Time-dependent functions for *b_max_*and *FP_bg_* were used to obtain a global, measurement of the HSPB8-TDP43 LC affinity represented by the complete FP-aggregation assay. The optimized time-dependent functions are also shown in Fig. 2C. Both *b_max_* and *FP_bg_* required two-component functions while the *K_D_* value was constant. The complete analysis and time-dependent functions are described in the *Materials and Methods* section.

Supplemental Fig. S3B shows a yADH assay run using the fluorescently labeled HSPB8, revealing a delay of yADH aggregation like that observed for unlabeled HSPB8. Supplemental Fig. S3E–F show size exclusion chromatography (SEC) analysis of labeled HSPB8 revealing the protein is monodisperse with a hydrodynamic radius consistent with monomeric HSPB8. The result is consistent with SEC analysis of the unlabeled HSPB8 in Supplemental Fig. S3D. Together, these characterizations indicate that the labeling of HSPB8 with NHS-fluorescein does not interfere with chaperone activity or alter the oligomeric state.

The initial time-dependent increase in *b_max_* in Fig. 2C arises from an increasing population of TDP43 LC species capable of interacting with HSPB8. The quadratic decay after the maximum value at 238 min is due to a gradual loss of these species. In Fig. 2C, the background fluorescence (*FP_bg_)* curve begins high and decays, rapidly and nonlinearly at first and then linearly and relatively slowly. Although HSPB8 can form a variety of heterooligomers (54, 55), it is known to exist as a mixture of primarily monomer and homodimer in equilibrium (51–53). The initial, rapid decrease in *FP_bg_* is consistent with the presence of a small amount of larger, slowly reorienting, HSPB8 homodimers that dissociate and contribute to a pool of smaller, more quickly reorienting, HSPB8 monomers. The best-fit curve for *FP_bg_* shows the nonlinear decay of the background fluorescence equilibrates at 94 min. Data from 94 min and later were thus used to calculate an average *K_D_* of 0.41 ± 0.25 µM, which did not require time-dependent fitting, consistent with a steady-state interaction between HSPB8 and TDP43 LC. The linear decay for *FP_bg_* after 94 min is likely due to quenching of the fluorophore. Supplemental Fig. S3C shows a plot of FP signal from HSPB8 alone recorded under the same conditions, illustrating this background FP signal is largely independent of TDP43 LC. Fig. 2E shows the FP binding isotherm calculated from these parameters with the corresponding regions of uncertainty. Table 1 summarizes the fitting parameters obtained from the FP and ThT analyses.

**Table 1:**
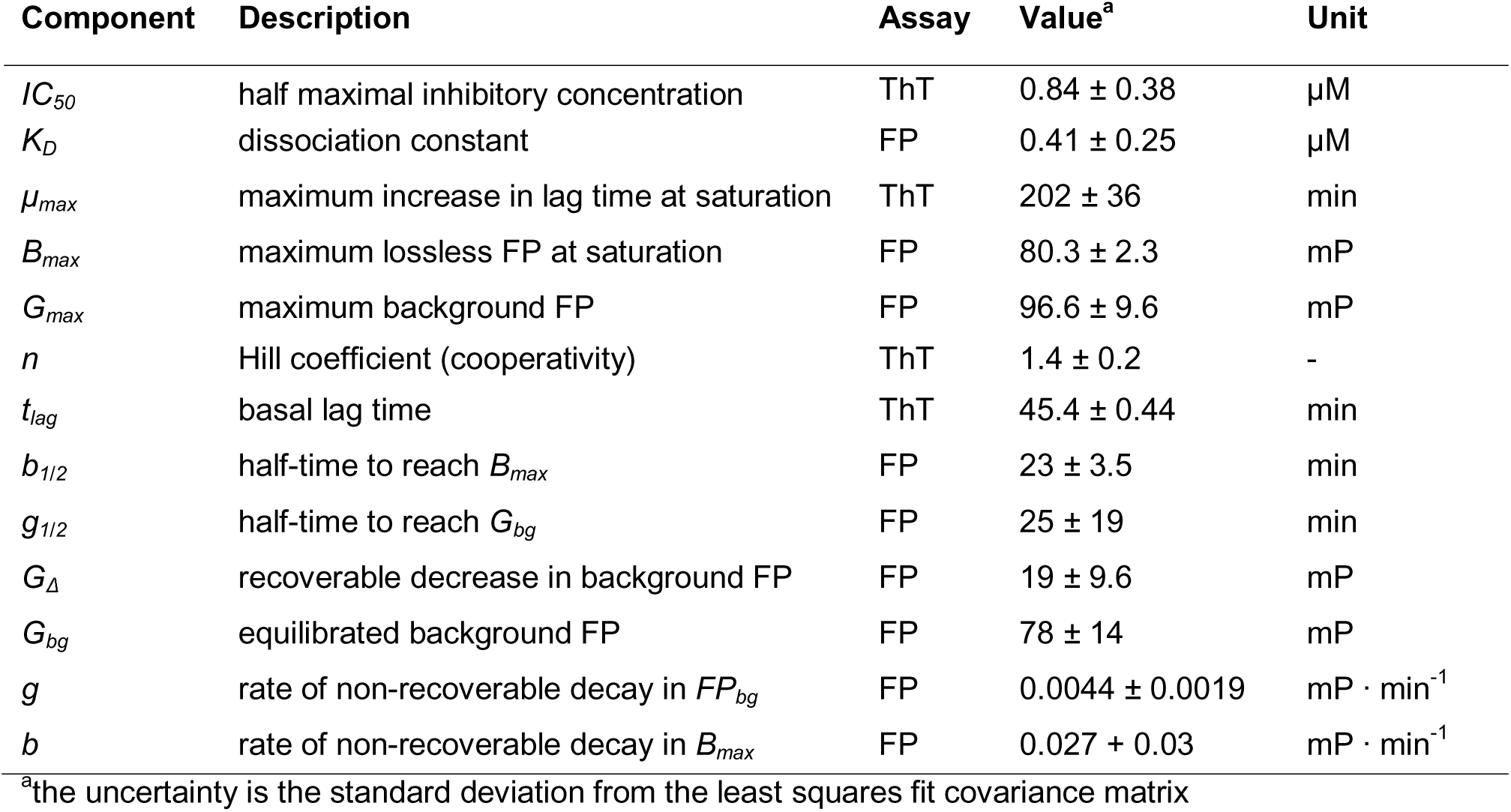
Parameters Derived from ThT and FP Assays.

| Component | Description | Assay | Value<sup>a</sup> | Unit |
| --- | --- | --- | --- | --- |
| $IC_{50}$ | half maximal inhibitory concentration | ThT | $0.84 \pm 0.38$ | $\mu\text{M}$ |
| $K_D$ | dissociation constant | FP | $0.41 \pm 0.25$ | $\mu\text{M}$ |
| $\mu_{max}$ | maximum increase in lag time at saturation | ThT | $202 \pm 36$ | min |
| $B_{max}$ | maximum lossless FP at saturation | FP | $80.3 \pm 2.3$ | mP |
| $G_{max}$ | maximum background FP | FP | $96.6 \pm 9.6$ | mP |
| $n$ | Hill coefficient (cooperativity) | ThT | $1.4 \pm 0.2$ | - |
| $t_{lag}$ | basal lag time | ThT | $45.4 \pm 0.44$ | min |
| $b_{1/2}$ | half-time to reach $B_{max}$ | FP | $23 \pm 3.5$ | min |
| $g_{1/2}$ | half-time to reach $G_{bg}$ | FP | $25 \pm 19$ | min |
| $G_{\Delta}$ | recoverable decrease in background FP | FP | $19 \pm 9.6$ | mP |
| $G_{bg}$ | equilibrated background FP | FP | $78 \pm 14$ | mP |
| $g$ | rate of non-recoverable decay in $FP_{bg}$ | FP | $0.0044 \pm 0.0019$ | $\text{mP} \cdot \text{min}^{-1}$ |
| $b$ | rate of non-recoverable decay in $B_{max}$ | FP | $0.027 \pm 0.03$ | $\text{mP} \cdot \text{min}^{-1}$ |

<sup>a</sup>the uncertainty is the standard deviation from the least squares fit covariance matrix

The FP (i.e. *b_max_*) and ThT data together provide more insight into the TDP43 LC aggregation process. The *b_max_*curve and ThT data from a concurrently performed ThT aggregation assay with identical starting material were normalized and plotted together in Fig. 2E. An initial increase in *b_max_* without significant ThT signal is consistent with HSPB8 interacting with pre-fibrillar TDP43 LC species, likely fibril-competent nuclei. A critical concentration of these nuclei is reached at *t_lag_* (t = 45 min), after which fibril formation and elongation occur, with ThT and *b_max_* fluorescence both increasing until the maximal nuclei concentration, the *b_max_* maximum, is reached at ∼238 min. Fibrils then become the dominant species, ThT fluorescence plateaus due to saturated binding sites on the fibrils or the depletion of free ThT, and *b_max_* trends downwards as the remaining TDP43 LC molecules are slowly converted to fibrils at the expense of nuclei bound to HSPB8.

To provide a visual characterization of the TDP43 LC during the aggregation process, negative stain TEM images were recorded at various timepoints along the aggregation curve. The images in Supplemental Fig. 4 show that initially the TDP43 LC samples are devoid of regularly shaped proteinaceous aggregates, oligomeric assemblies, and fibrils. Short fibrils then form in conjunction with an increase in ThT fluorescence. Elongated, bundled fibrils are observed in the images once the ThT fluorescence plateau is reached. Across all time points, we did not observe any well-defined assemblies attributable to HSPB8-TDP43 LC complexes or TDP43 LC nuclei. If present, these assemblies were too small for the resolution of the TEM experiment. Small, white, puncta are observed in some of the images, but they are not regularly shaped or visually identical across the images.

### HSPB8 Reduces Diffusion within TDP43 LC Droplets

Previous work has shown that HSPB8 plays a role in regulating liquid droplets (30) and stress granules (26). We explored the interaction of HSPB8 with TDP43 LC liquid droplets with fluorescence recovery after photobleaching (FRAP) experiments. Supplemental Fig. S5 shows a diagram of the sample setup for the confocal microscopy experiments. Supplemental Fig. S5 also shows TDP43 LC liquid droplets are reliably produced by dilution into PEG-8000 with and without HSPB8. The confocal images in Fig. 3B and Supplemental Fig. S5 show that HSPB8 preferentially localizes to TDP43 LC droplets. The shallow FRAP curves in Fig. 3A and the persistence of a distinct bleached spot in Fig. 3B indicate that HSPB8 and TDP43 LC co-colocalize and have limited mobility. There is, however, a small but significant increase in the maximum fluorescence recovered after 60 s for TDP43 LC relative to HSPB8 (Welch’s t-test, p = 0.016, 95% confidence). Additionally, based on the τ_1/2_ recovery parameters shown in Table 2, TDP43 LC fluorescence recovers two times faster than HSPB8 fluorescence in the mixed droplets. Although droplets of TDP43 LC and HSPB8 at the surface of the microscope slide lack fluidity in the FRAP measurements, Supplemental Fig. S6 shows droplet fusion in the bulk solution. Unfortunately, the rapidly moving droplets in solution could not be bleached. Conversely, droplets formed by TDP43 LC alone display significant fluidity at the surface of the microscope slide, with the fluorescence signal shown in Fig. 3C and 3D recovering completely after only 30 s. Fig. 3C and 3D show recovery for whole-droplet bleaching, indicating TDP43 molecules exchange between the concentrated and dilute phases. Fitting parameters from all FRAP assays are shown in Table 2. Altogether, the FRAP data are consistent with HSPB8 reducing TDP43 LC mobility in liquid droplets.

**Figure 3.**
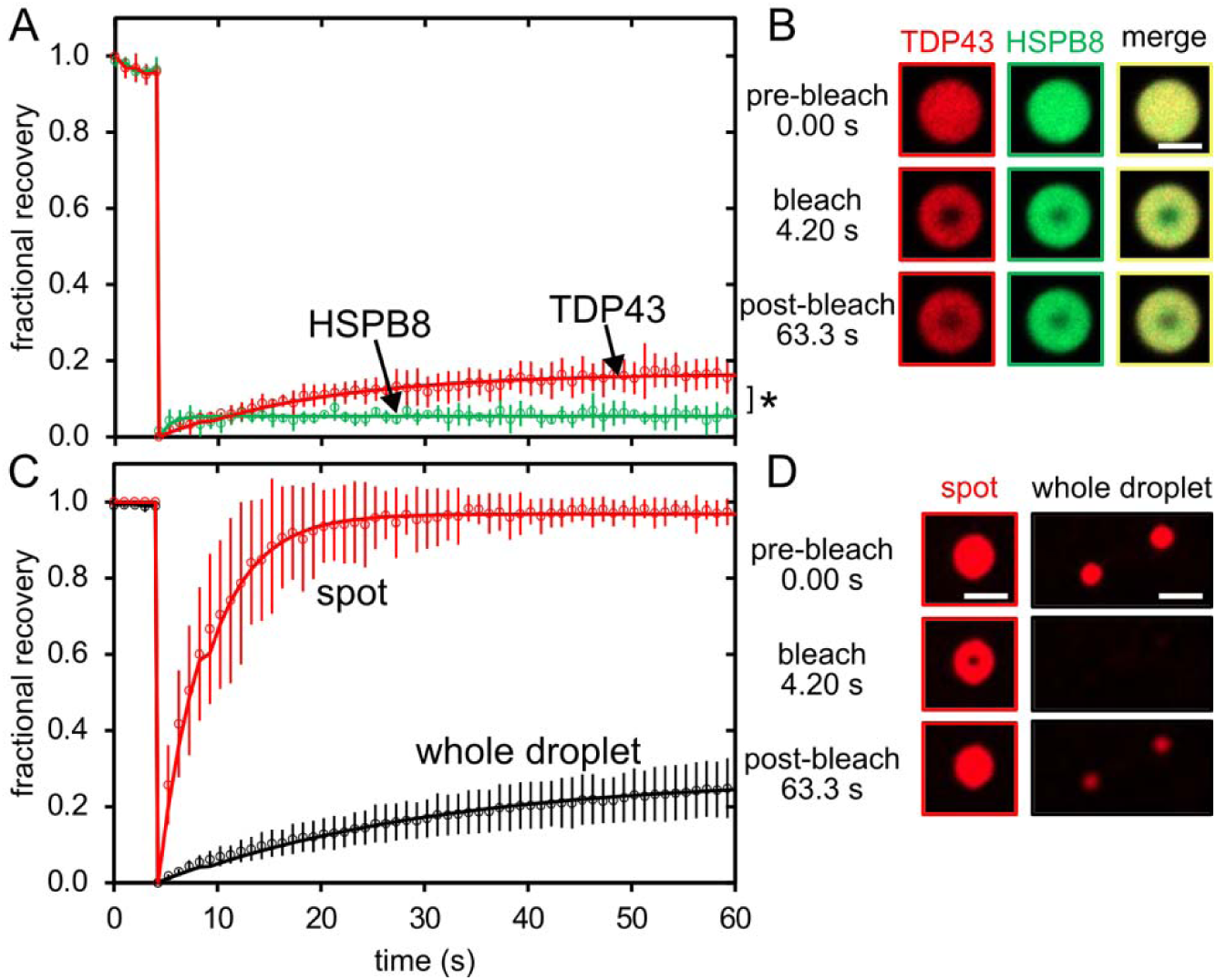
TDP43 LC liquid droplets are less fluid in the presence of HSPB8. (A) Fluorescence recovery after photobleaching (FRAP) curves recorded on liquid droplets containing both fluorescein-HSPB8 (Green) and Alexa Fluor 647-TDP43 LC (Red). (B) Confocal images of a representative liquid droplet from the FRAP measurements shown in panel (A). (C) FRAP curves for Alexa Fluor 647-TDP43 LC liquid droplets alone. The red curve represents a spot based FRAP measurement, the black curve represents a whole droplet bleaching measurement. (D) Confocal images of representative liquid droplets from the FRAP measurements shown in panel (C). The white inset scale bars are all 2 μM. Each FRAP datapoint is the average of n = 4 measurements and the error bars represent ±1 standard deviation. For bleaching subregions of droplets, the area and laser intensities were the same. Whole droplets were bleached using the entire area of the droplet.

**Table 2:** Parameters Derived from the FRAP Assays^a^.

| Component | HSPB8<br>(in TDP43 LC droplets) | TDP43 LC (with<br>HSPB8) | TDP43 LC<br>(alone) | TDP43 LC<br>Whole Droplet |
| --- | --- | --- | --- | --- |
| immobile fraction ($\alpha$) | $0.9456 \pm 0.001$ | $0.8332 \pm 0.002$ | $0.0323 \pm 0.002$ | $0.7223 \pm 0.004$ |
| $\tau_{1/2}$ | $4.98 \pm 0.26$ | $14.78 \pm 0.46$ | $7.26 \pm 0.05$ | $22.46 \pm 0.62$ |

<sup>a</sup>the uncertainties are the standard deviation from the least squares fit covariance matrix

### HSPB8 ACD Domain Interacts with the Second Core Region of TDP43 LC

Previous studies on sHSP have suggested that the α-crystallin domain (ACD), the disordered N-terminal region (NTR), or both might play a role in chaperone activity (56–61). To identify the regions involved in the interaction between TDP43 LC and HSPB8, we used cross-linking mass spectrometry (XL-MS), a technique that has been used previously to assess the region of interaction between FUS and HSPB8 in liquid droplets (30).

HSPB8 and TDP43 LC were studied independently and jointly to optimize, identify, and categorize the crosslinked species by SDS-PAGE. Fig. 4A and 4B show homomeric crosslinking for both TDP43 LC and HSPB8, consistent with the propensity of TDP43 LC to assemble at neutral pH (45) and HSPB8 to form a dimer (52). Interprotein crosslinking with TDP43 LC is especially challenging due to its namesake low complexity amino acid composition. Supplemental Fig. S7 shows that including both the N- and C-termini and the overhang after purification tag cleavage, TDP43 LC has only three basic and five acidic residues available for crosslinking. Conversely, HSPB8 has 37 basic and acidic residues available for crosslinking with basic residues biased to within the ACD and acidic residues spread across the entire sequence.

**Figure 4.**
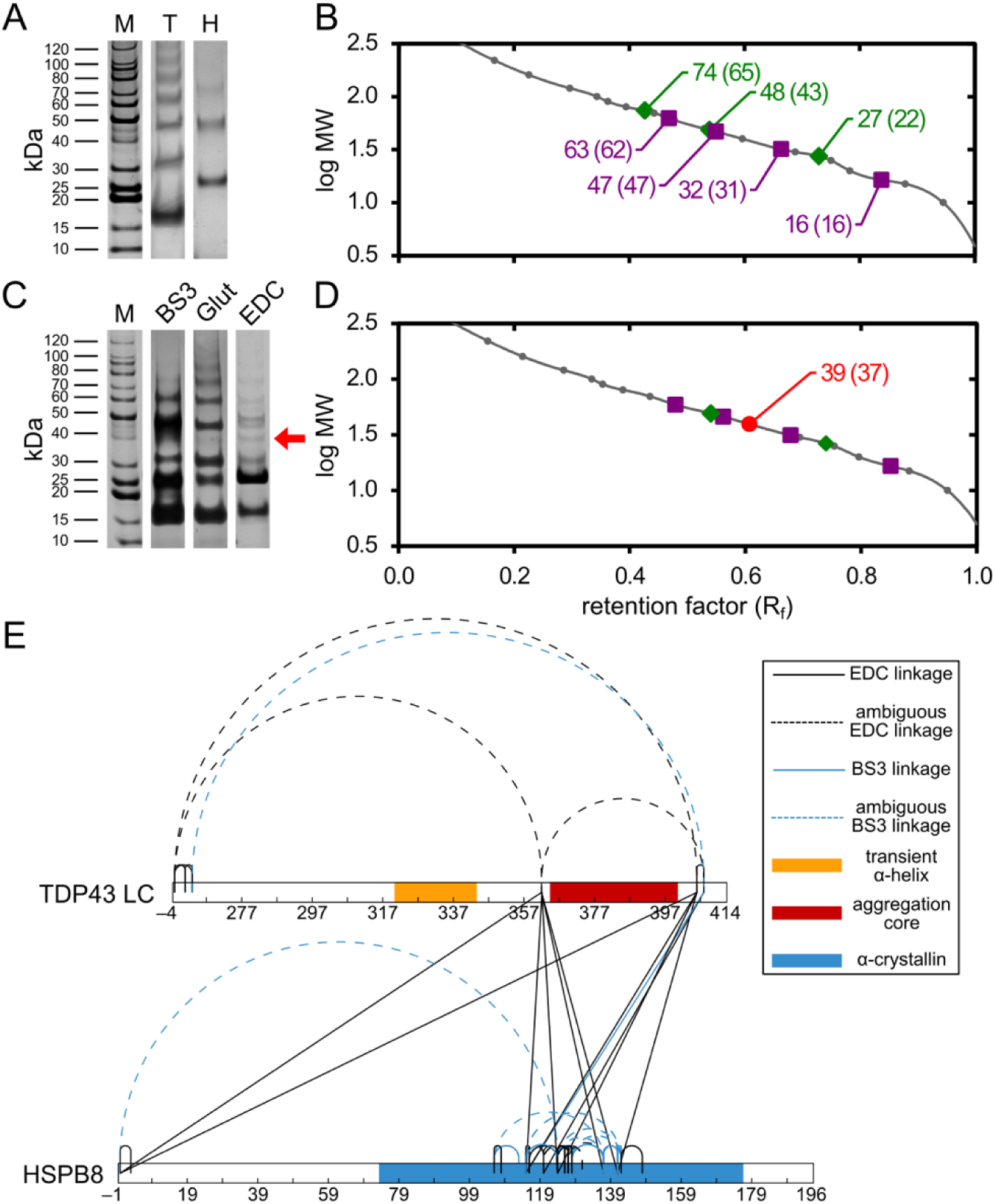
Crosslinking paired with mass spectrometry (XL-MS) identifies a region of interaction between HSPB8 and TDP43 LC. (A) Strips from an SDS-PAGE gel for the molecular weight marker (M), TDP43 LC only (T), and HSPB8 (H) only. Labels to the left of M correspond to the molecular weights of the marker bands. Both T and H are crosslinked with glutaraldehyde. All gels are stained with Coomassie blue, T was also silver stained. (B) Plot of the best-fit cubic spline function (solid gray line) to the molecular weight marker bands (gra circles) from the SDS-PAGE gel in (A) used to calibrate migration through the gel to molecular weight. Purple squares correspond to T and green diamonds correspond to H from panel (A). Only the first 4 bands (monomer, dimer, trimer, and tetramer) are plotted for TDP43 LC and only the first 3 bands are plotted for HSPB8 (monomer, dimer, trimer). Labels indicate the predicted molecular weight from the best-fit cubic spline function, with the value expected from the oligomeric state and primary sequence in parenthesis. (C) Strips from an SDS-PAGE gel for the molecular weight marker (M) and three separate mixtures of TDP43 LC and HSPB8 crosslinked by either BS3, glutaraldehyde (Glut), or EDC. Labels to the left of M correspond to the molecular weights of the marker bands. All gels are stained with Coomassie blue, BS3 and Glut were also silver stained. The red arrow points to the location of a TDP43 LC and HSPB8 heteromeric crosslink. (D) Plot of the best-fit cubic spline function (solid gray line) to the molecular weight marker bands (gray circles) in M from (C) used to calibrate migration through the gel to molecular weight. The purple squares and green diamonds correspond to the identified HSPB8 and TDP43 LC species in panels (A & B) that are also present in the EDC strip from panel (C). The red circle indicates the position of the band with the red arrow in (C), labeled with the molecular weight predicted by the calibration plot and with the expected value for a 1:1 HSPB8-TDP43 LC complex in parenthesis. (E) Crosslink map based on inter- and intrapeptide linkages identified by XL-MS from BS3 and EDC crosslinking. Interpeptide loops are shown as unambiguous, intrapeptide loops are shown as ambiguous, and the domains are defined as transient alpha helix, residues 321–343; aggregation core, residues 365–400; and the alpha crystalin domain (ACD), residues 74–176. The crosslinking map is a modified map generated by xiNET (92).

Comparison of the SDS-PAGE images in Fig. 4A with 4C and the cubic spline analyses in Fig. 4B with 4D shows that the amine-amine linkers, bis(sulfosuccinimidyl)suberate (BS3) and glutaraldehyde (Glut), were unable to produce a distinct HSPB8-TDP43 LC linkage. On the contrary, the data in Figure 4 show that the acid-amine linker, 1-ethyl-3-(3-dimethylaminopropyl)carbodiimide (EDC) paired with N-hydroxysulfosuccinimide (sNHS) proved to be highly successful in linking a TDP43 LC monomer with an HSPB8 monomer, the only condition tested which produced a novel crosslinked band. The EDC SDS-PAGE gel strip is only stained with Coomassie blue, while BS3 and Glut gels are both double-stained with Coomassie blue and Silver Stain. Silver stain was used to rule out sparsely populated states because the limit of detection (LOD) for Coomassie blue is ∼0.1–0.5 ug (62) while silver stain is approximately 400–2,000X more sensitive with an LOD of 0.25 ng (63). Furthermore, BS3 has a linker length of 11.4 Å increasing the flexibility and likelihood of a crosslink, while EDC is a 0 Å crosslinker, as the final bond formed is spacer-free native-like amide bond, further highlighting the close HSPB8-TDP43 LC contact indicated by the observed EDC cross-linkage.

To obtain sequence-specific interaction sites, the crosslinked proteins were enzymatically digested, and the resultant peptides analyzed by XL-MS. Fig. 4E shows a cartoon summarizing the XL-MS analysis and Supplemental Tables S1–S4 list the observed crosslinked peptides. The ambiguous linkages cannot be definitively assigned as occurring between two of the same protein or within the same protein without additional samples and analysis. However, 22 of the 26 (or 85%) unique HSPB8-HSPB8 crosslinked sites are unambiguous linkages within the ACD of HSPB8. In addition, the only confirmed dimer linkage for HSPB8 was found on the ACD, suggesting the dimer is formed through the ACD.

The identified hetero-interprotein crosslinks provide evidence for interactions between the NTR (residues 1–73) and ACD of HSPB8 (residues 74–176) with a fibril-forming core of TDP43 LC (residues 365–400), previously identified by Fonda et al. (24). This result substantiates previous work showing that the both the NTR and ACD of HSPB8 may play roles in substrate recognition, binding, and activity (56–61, 64).

## Discussion

sHSP are essential molecular chaperones that help preserve cellular proteostasis, uniquely capable of recognizing and stabilizing misfolded or aggregating proteins through remarkably low affinity, transient PPIs. Quantifying the dissociation constants of protein chaperones is incredibly challenging due to inherently weak binding affinities of these PPI, a challenge compounded by the short-lived nature of their substrates. These substrates are highly sensitive, prone to a cascade of misfolding, oligomeric aggregation, and subsequent precipitation, making them difficult to capture. In this manuscript, we have demonstrated two methods to measure the affinity of HSPB8 for a suspected fibril nucleating species of the TDP43 LC. Indirect measurements (IC_50_) were achieved by assessing how HSPB8 alters the aggregation kinetics of TDP43 LC in a ThT aggregation assay (Fig. 1), while direct measurements (K_D_) were obtained through time-dependent fluorescence polarization (Fig. 2). Both methods effectively yield dissociation constants (IC_50_ vs K_D_) that are also in good agreement (ThT, IC_50_ = 0.84 ± 0.36 µM; FP, K_D_ = 0.41 ± 0.25 µM). Due to the simple environment of our in vitro assay, rough equivalence of the measured IC_50_ value with a true K_D_ is expected. In doing so we have also gained insight into the mechanism of interaction, where HSPB8 delays TDP43 aggregation by inhibition of primary nucleation. A similar chaperoning mechanism has been observed for DNAJB6, a member of the HSP40 family (65).

Although many sHSP readily form higher order oligomers (66–69), these forms are thought to act as “storage” until dissociation into dimers or monomers enables chaperone activity (69–71). Our results are consistent with monomeric HSPB8 binding to monomeric TDP43 LC. This conclusion is supported by several observations. First, the lack of cooperativity (Fig. 1; Hill coefficient, 1.4 ± 0.2) in binding indicates TDP43 LC binds only as a monomer. Second both the background FP and overall change in FP is too low to corroborate either a dimer of HSPB8 interacting with a TDP43 LC monomer or a HSPB8 PPI with anything more than a monomer of TDP43 LC (Fig. 2). Finally, the crosslinking SDS-PAGE gels do not show any evidence of dimeric PPI between HSPB8 and TDP43 (Fig. 4). These results, however, do not preclude the existence of a HSPB8 oligomer. On the contrary, while we observe no evidence for high order oligomers, we observed evidence for a small population of dimeric HSPB8 in equilibrium with monomeric HSPB8. This equilibrium is verified by the MALDI-TOF of HSPB8 presenting a low intensity dimer peak (Supplemental Fig S8), the transiently high background FP likely arising from a small population of dimeric HSPB8 dissociating at higher temperatures (Fig. 2C), and the crosslinking gels consistently depicting both a monomer and dimer for HSPB8 (Fig. 4A and 4C). In addition, based on the high levels of inter-peptide crosslinking within the ACD determined by XL-MS (Fig. 4C and Supplemental Tables S1–S4), we suspect that this region is likely involved in HSPB8 self-association. In other sHSP, the ACD is known to play a significant role in lower level oligomerization, such as dimerization (72–74).

Our data also suggest that TDP43 LC nuclei are in equilibrium with fibrils and that these nuclei are the substrate recognized by HSPB8. The change in FP observed here is not drastic enough to indicate binding to fibrils, and even after 16 h, there still exists concentration-dependent FP, thus some amount of fibril forming competent TDP43 LC nuclei must be available for HSPB8 to bind, even after the system has reached equilibrium in the ThT assay reporting on fibril formation. In addition, HSPB8 only increases the *t_lag_* and does not alter the rate of fibril elongation, *k*_app_, supporting that HSPB8 only interacts with TDP43 LC nuclei. Changes in *k*_app_ would suggest a change in the fibril elongation process. Finally, TEM analysis showed no clear signs of HSPB8 interference in the eventual formation of fibrils or direct complex with fibrils through PPI (Supplemental Fig. S4). TDP43 LC nuclei were also invisible to TEM, possibly a result of their existence as monomers.

Existing literature on liquid droplets formed by LC proteins often show an increase in fluidity and dynamics with the addition of an sHSP or chaperone. For example, partitioning of HSPB8 into droplets formed by full-length FUS protein maintains droplet fluidity and is driven by HSPB8 interactions with the FUS LC domain (30). Similarly, HSPB1 binds directly to the TDP43 RNA-binding and LC domains, delaying fibril formation and facilitating droplet disassembly (75). Our FRAP experiments reveal a contrary case where the presence of HSPB8 reduces the molecular motion in a liquid droplet (Fig. 3). By confining and reducing the mobility of TDP43 LC within the droplets, the cell’s degradation pathways may more efficiently recognize and process aggregating or misfolded proteins (35, 76–79). The holdase mechanism of HSPB8 may itself be involved with the sequestering process, not only limiting the diffusion and conformational mobility of the TDP43 LC, but also preventing exchange with the bulk solution. However, in our experiments, FRAP of liquid droplets is inherently limited to immobilized droplets at the microscope slide surface, potentially causing a self-selection bias for the least mobile population of droplets. Droplet fusion events are observed above the microscope slide surface like those shown in Supplemental Fig S5, indicating at least some fraction of the TDP43 LC and HSPB8 containing droplets exhibit very fluid-like properties.

Our crosslinking results implicating a HSPB8-TDP43 LC chaperone interaction after residue 367 in TDP43 LC seem to contradict recent structural studies on tissue from ALS patients with type B FTLD-TDP (residues 282–360) (38) and type A FTLD-TDP (residues 272–360) (39). In the context of an earlier study showing a transient helix formed by residues 321–340 facilitating TDP43 LC self-association and liquid droplet formation (41), these ex vivo structural models suggest TDP43 LC fibril formation arises from the conversion of a transient alpha helix into a beta strand involving residues clearly preceding the HSPB8-TDP43 LC interaction site we identified. However, in vitro results contend that both an expanded alpha helical region (residues 311–360) and a second fibril forming region (residues 365–400) form amyloid fibrils (14, 24, 80, 81). Furthermore, the same study showing that TDP43 LC self-association is mediated by interactions between the alpha helix also showed that residues 382–385 in the second fibril-forming core participate in the self-association of TDP43 LC (41).

One interpretation of these combined results is a model in which TDP43 LC nuclei form through misfolding-induced self-association of residues in the second fibril-forming core (residues 365–400). Pathological aggregation then proceeds via fibril formation involving the first fibril forming core (residues prior to 360) due to high local concentrations of TDP43 LC. In this light, HSPB8 could delay fibril formation through holdase activity by inhibiting misfolded TDP43 LC self-association via interactions in the second fibril-forming core. Fig. 5 illustrates the HSPB8-TDP43 LC model described here. Disease-specific mechanisms promoting fibril formation such as amino acid changes in the TDP43 LC helical region would not be intercepted by the HSPB8 chaperone system under this model. Furthermore, mutations disrupting the HSPB8-TDP43 LC interaction in the second fibril-forming core region would also be detrimental, which is consistent with the 16 disease-related mutations between residues 362–406 (82).

**Figure 5.**
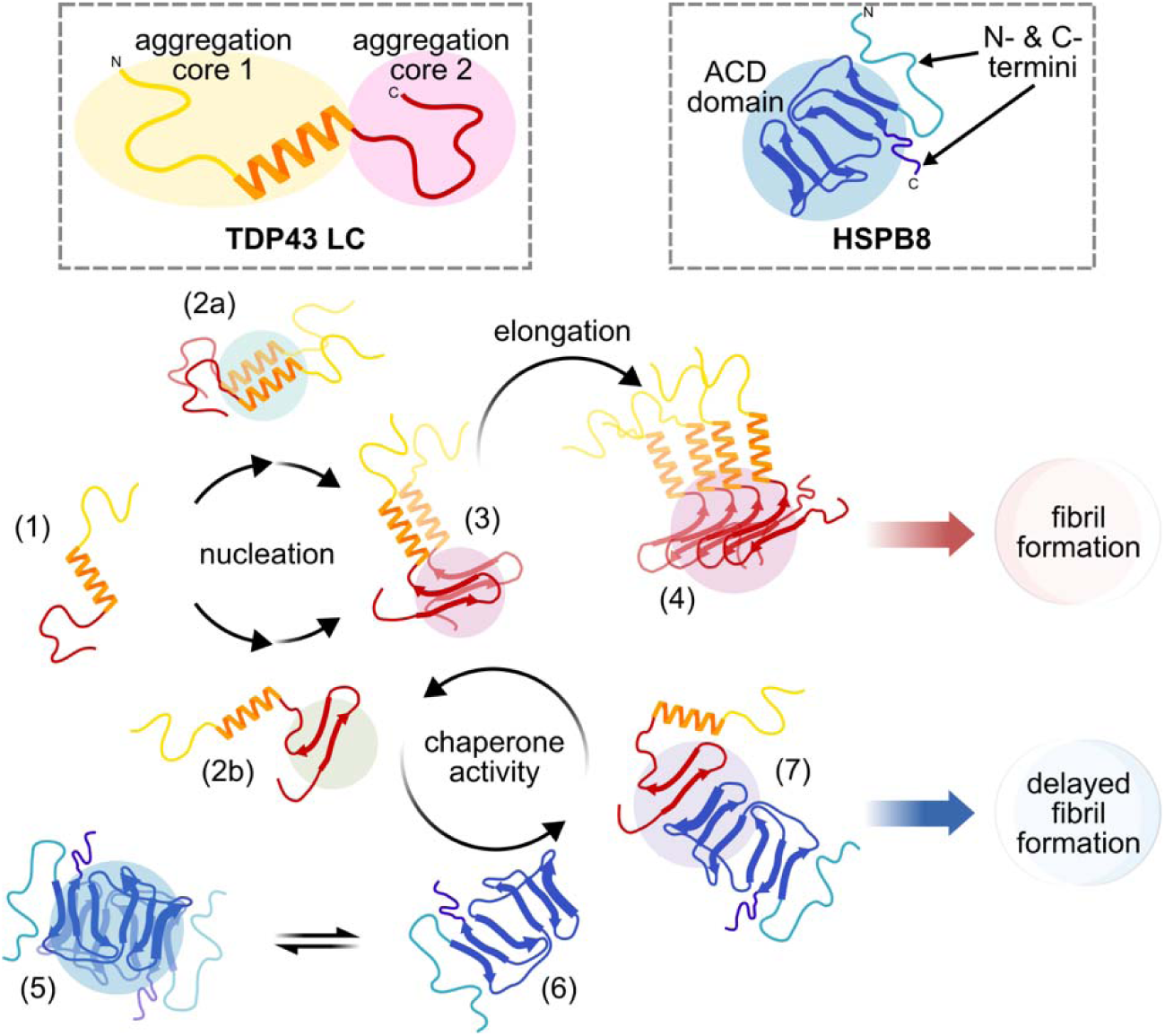
Putative mechanism of TDP43 LC aggregation inhibition by HSPB8. Monomeric TDP43 LC (1) undergoes a nucleation process, oligomerizing (2a) and misfolding (2b) or both. These nuclei (oligomers or misfolded species) promote the misfolding and/or binding of additional misfolded TDP43 LC (3) which then begins the elongation and amyloid fibril formation process (4). Dimeric HSPB8 (5) is in equilibrium with the monomer (6) which disrupts the nucleation process by binding a misfolded nuclei (2b) or favors binding a monomeric species (3). An interaction between the ACD of HSPB8 and the aggregation core of TDP43 LC forms a transient complex (7). Aggregation is delayed through HSPB8 holdase activity (26, 93). The domain coloring scheme is the same as Fig. 4E with the transient TDP43 LC α-helix in orange, the TDP43 LC aggregation core in red, and the HSPB8 α-crystallin domain in blue.

However, while we can rule out HSPB8 interactions with the extreme N-terminus of TDP43 LC, we note that the residues able to crosslink are simply not sufficiently distributed to capture the full extent of the HSPB8-TDP43 LC interaction. It is entirely possible that HSPB8 recognizes both fibril forming regions of TDP43 LC, but an interaction with the alpha helical region (residues 311–360) is unobserved due to a lack of residues able to crosslink in the region. It should also be noted that the results presented here pertain to only the C-terminal TDP43 low complexity domain and do not report on any potential interactions between HSPB8 and the TDP43 N-terminus and RNA binding domains. Regardless, the work presented here is relevant to TDP43 aggregation in disease, as segments of the LC domain form the pathological aggregates in brain tissue for FTD patients (38, 39), and C-terminal truncations (TDP25) are associated with TDP43 pathology (37).

In conclusion, our study provides valuable insights into the specific interactions and mechanisms by which HSPB8 modulates TDP43 LC aggregation. Understanding these interactions at a molecular level can inform future therapeutic strategies aimed at mitigating protein aggregation in neurodegenerative diseases. In the biological milieu, HSPB8 is known to form complexes with several different sHSP, ATP-dependent chaperones, and other proteins (26, 55, 83–85). Additionally, HSPB8 has several known locations of phosphorylation, which has been shown to alter the activity of HSPB8 and other sHSP (86–88). These factors likely influence HSPB8 chaperone activity in vivo, placing a limitation on the interpretations and extrapolations of these in vitro results. To fully appreciate the broader implications of the interactions observed here in disease pathology, further research is needed to explore the complex interplay between HSPB8 and its substrates through higher resolution structural studies and in cellular contexts.

## Materials and Methods

### TDP43 LC Growth and Expression

WT TDP43 LC (residues P262–M414) were recombinantly produced using a pHIS-parallel plasmid (89) that includes a TEV-cleavable N-terminal 6xHis-tag. The plasmids were transformed by heat shock into chemically competent BL21(DE3) E. coli cells. All cells were grown in a shaker incubator at 37 °C and 220 revolutions per minute (RPM). Bacterial cultures were grown in Luria Broth media supplemented with 1% (w/v) glucose in the presence of 100 μg/ml ampicillin to an optical density at 600 nm of 0.6–1.0 measured using a 1 cm pathlength cuvette on an Eppendorf Biospec Kinetic before adding isopropyl β-D-thiogalactopyranoside (IPTG) to 0.5 mM to induce protein expression. The cultures were grown for another 3 h before harvesting the cells by centrifugation at 6,000 g for 15 min. The cell pellets were flash frozen in liquid nitrogen and stored at −80 °C until purification.

### TDP43 LC IMAC Purification

Immobilized metal affinity chromatography (IMAC) purification of the TDP43 LC was done as reported previously by Fonda et al. (24) In brief, the cell pellet was resuspended in 35 mL of 6.0 M guanidinium chloride (Gu-HCl), 1% v/v Triton X-100, 500 mM sodium chloride, 50 mM 2-amino-2-(hydroxymethyl)propane-1,3-diol hydrochloride (Tris-HCl), pH 7.5 containing 3 pellets of mini EDTA-free Pierce Protease Inhibitor (ThermoFisher Scientific). The resuspended pellet was sonicated at a 12% duty cycle for 20 min at 30% power using a Branson SFX-250 Sonifier with a 1/4 in. microtip. The lysate was centrifuged at 75,600 g for 30 min at 4 °C, and the supernatant decanted, and filtered through a 0.22 µm PES syringe filter (Millipore). IMAC was performed using a Bio-Rad NGC Quest 10 Plus chromatography system and a 5 mL Bio-Scale^TM^ Mini Nuvia^TM^ IMAC column (Bio-Rad). The column was equilibrated with 6.0 M Urea, 500 mM sodium chloride, 20 mM 2-[4-(2-hydroxyethyl)piperazin-1-yl]ethanesulfonic acid (HEPES), pH 7.5 before loading the filtered cell supernatant. The column was then washed with the equilibration buffer containing 20 mM imidazole, and then the protein was eluted isocratically by raising the imidazole concentration to 200 mM. The presence of highly pure TDP43 LC was assessed by non-reducing denaturing SDS-PAGE.

### TDP43 LC His-tag Removal

IMAC purified TDP43 LC was diluted to 20 µM with buffer containing 0.5 M Gu-HCl, 1 mM 1,4-bis(sulfanyl)butane-2,3-diol (DTT), 20 mM 2-morpholin-4-ylethanesulfonic acid (MES), pH 6.0. Tobacco Etch Virus protease (TEV, prepared in-house) was then added to a final concentration of 0.2 µM. The cleavage reaction was run at room temperature (∼20 °C) under quiescent conditions. After 4 and 24 h the cleavage reaction mixture was recharged with an additional 1 mM DTT and 0.2 µM TEV, giving final concentrations of 3 mM DTT and 0.6 µM TEV. SDS-PAGE and MALDI-TOF were used to confirm that His-tag cleavage. The TEV reaction was mixed with 5 mL of Nuvia IMAC Ni-charged resin (Bio-Rad) in equilibration buffer and gently swirled before incubation for 30 min at 4 °C. His-cleaved TDP43 LC was eluted by pouring the column over a glass Econo-Column (Bio-Rad), collecting the flow through, and washing the resin with 10 column volumes of 6 M Urea, 500 mM sodium chloride, 15 mM Imidazole, and 20 mM Tris-HCl, pH 7.5. After TEV cleavage, non-native residues GAMD remain at the N-terminus of the protein.

### TDP43 LC Cation Exchange Chromatography (CEX) Purification

Eluent and wash fractions containing His-cleaved TDP43 LC were dialyzed into 6 M Urea, 10 mM 2-[4-(2-sulfopropyl)piperazin-1-yl]propylsulfonic acid (PIPPS), pH 4.0 using Spectra Por 3 regenerated cellulose, 29 mm flat width dialysis tubing. The protein was then syringe filtered (sterile 0.22 µm, PES) and loaded onto an UNO S6 Column (Bio-Rad) equilibrated in CEX buffer using a Bio-Rad NGC Quest 10 Plus Chromatography system. Purified protein was eluted using the same buffer with a 0–0.2 M sodium chloride gradient over 15 column volumes (CV). Elution fractions were assayed by non-reducing SDS-PAGE. Fractions of his-cleaved TDP43 LC with no impurities observed in the SDS-PAGE gel were pooled, aliquoted, flash frozen in liquid nitrogen, and stored at −80 °C.

### HSPB8 Growth and Expression

HSPB8 was recombinantly produced using a pQTEV plasmid (AddGene #34710) (90) containing a TEV cleavage site and an N-terminal 6xHis-tag. The plasmid was transformed into chemically competent BL21(DE3) E. coli cells by heat shock. Bacterial cells were cultured, induced, and harvested identically to the method described for TDP43 LC.

### HSPB8 IMAC Purification and His-tag Removal

The cell pellet was thawed on ice for 30 minutes, then resuspended in 36 mL of 25 mM Tris-HCl, pH 8.0 500 mM sodium chloride, 0.25 mg/mL lysozyme, and 3 pellets of mini EDTA-free Pierce Protease Inhibitor (ThermoFisher Scientific). The resuspended pellet was placed on ice and sonicated twice using a Branson SFX-250 Sonifier with a 1/4 in. microtip at 30% power, with a 12% duty cycle, for 10 min. The ice was replaced between each sonification. The lysate was centrifuged at 75,600 g at 4 °C for 30 minutes, then the supernatant was decanted and filtered through a 0.22 µm PES syringe filter (Millipore).

IMAC was completed on a Bio-Rad NGC Quest 10 Plus Chromatography system using a 5 mL Bio-Scale^TM^ Mini Nuvia^TM^ IMAC column (Bio-Rad). The column was pre-equilibrated with 25 mM Tris-HCl, pH 8.0, 500 mM sodium chloride, and 5 mM DTT. DTT was added to buffers immediately before use. The supernatant was loaded onto the column and washed with the equilibration buffer containing 20 mM imidazole. HSPB8 was eluted by the same buffer with an imidazole gradient of 20–500 mM. Elution fractions were assessed by SDS-PAGE then fractions without visible impurity bands were combined and diluted to 15 µM HSPB8 with equilibration buffer.

TEV protease (prepared in-house) was added immediately after dilution at a molar ratio of 100:1 (HSPB8:TEV) and transferred to Spectra Por 3 regenerated cellulose, flat width 29 mm, dialysis tubing. The TEV reaction was dialyzed at 4 °C for 18–20 hours against 100 mL of 25 mM Tris-HCl, pH 8.0, 5 mM DTT for every 1 mL of 15 µM HSPB8. The protein was harvested and diluted with an equal volume of 25 mM Tris-HCl, pH 8.0, 500 mM Sodium chloride, 20 mM imidazole then 5 mL of Bio-Rad Nuvia IMAC Ni-charged resin in the same diluent buffer was added to the crude cleavage reaction. The mixture was gently swirled before incubation for 30 min at 4 °C with gentle rotation. His-cleaved HSPB8 was eluted by pouring the mixture over a glass Econo-Column (Bio-Rad), collecting the flow through, and washing the resin with the diluent buffer. Eluent and wash fractions containing HSPB8 were then dialyzed overnight in 25 mM Tris-HCl, pH 8.0, and 5 mM DTT. Protein was placed at 4 °C for short term storage (1 week) or combined with an equal volume of 50% V/V glycerol and kept at −80 °C for long-term storage.

### Thioflavin T Aggregation Assays

A stock ThT solution, >500 μM, was prepared by dissolving solid ThT (Acros Organics) in ultrapure water. The stock solution was filtered through a sterile 0.22 μm PES membrane (Millipore). The ThT concentration was determined by absorbance at 412 nm on an Eppendorf BioSpectrometer Kinetic using a molar absorption coefficient, ε_412_, of 36,000 M^-1^ cm^-1^ and a 1 mm path length cuvette. Working solutions were prepared by diluting the stock solution to 60 µM ThT, 450 mM sodium chloride, and 20 mM Tris-HCl, pH 7.5. All ThT solutions were encased in aluminum foil and stored at 4 °C for up to 7 d.

CEX purified TDP43 LC was removed from the −80 °C and thawed at 4 °C. The TDP43 LC was then concentrated to 600 µM using an Amicon Ultra 0.5 mL 3K MWCO centrifugal filter and centrifugation at 14,000 g in 5–10 min intervals. TDP43 LC was diluted to 60 μM with 20 mM Tris-HCl, pH 7.5 immediately before use. HSPB8 was concentrated to 60 µM into 20 mM Tris-HCl, pH 7.5, 5 mM DTT using Amicon Ultra 0.5 mL 3K MWCO centrifugal filters and centrifugation at 5,000 g in 5–10 min intervals at 4 °C. At each interval in the protein concentration process, the tubes were inverted to ensure the most uniform concentration of the protein possible.

To help reduce temperature edge effects, the outer wells of a black-bottom 96 well plate were filled with 200 μL ultrapure water each, sealed with low autofluorescence Thermo Scientific™ Nunc™ adhesive seals, and preheated to 37 °C. Each sample well was adjacent to at least 2 water wells or 2 sample wells in every direction.

Finally, 300 µL solutions of each condition for the aggregation assay were prepared from stock solutions of 60 µM ThT, 60 µM TDP43, a varying amount of HSPB8, and 20 mM Tris-HCl. The final buffer components are 20 mM Tris-HCl, pH 7.5, 150 mM sodium chloride, 1.7 mM DTT, with 20 µM ThT, 20 µM TDP43 LC, and variable amounts of HSPB8. Each well was then filled with 90 µL of sample, immediately sealed with low autofluorescence Thermo Scientific™ Nunc™ adhesive seals, and placed inside a BioTEK Synergy H1 plate reader. ThT fluorescence measurements were recorded in 5 min intervals for up to 18 h, at 37 ° C, with excitation at 440 nm, emission at 485 nm, and 30 s of linear shaking at the instrument’s maximum speed before each read. The bandwidth was 9 mm for both excitation and emission. The instrument gain was fixed at 100 and an optimal read height of 5.75 mm was determined by the instrument software for the 90 µL samples.

### Thioflavin T Aggregation Data Analysis

Data analysis was performed as described by Malmos et al. (46) In brief, data were normalized using:

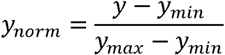

where *y_norm_* is the normalized value, y is the raw fluorescence intensity, *y_min_* and *y_max_* are the minimum and maximum fluorescence values for a single trial. Using non-linear least squares analysis in Python the normalized data was fit to the following equation:

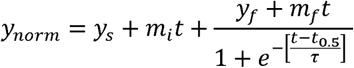

where *t* is time. The remaining parameters were obtained from the least squares analysis, where *y_s_* and *y_f_* are the initial and final ThT fluorescence intensities, *m_i_* and *m_f_* are the baseline slopes before and after the rapid rise in fluorescence, *t_0.5_* is the time to achieve half of the maximal fluorescence intensity, and τ is the elongation time constant.

The apparent fibril elongation rate*, k_app_*, and the lag time to fibril formation, *t_lag_*, were obtained from:

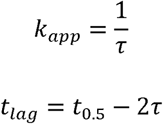

The final plots after fitting shown in Fig. 1A were normalized for ease of comparison.

Using non-linear least squares analysis in Python the lag time plot as a function of HSPB8 concentration was fit to a modified Hill Equation:

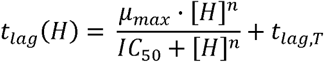

where *µ_max_* is the maximum achievable lag time, *t_lag,T_* is the basal lag time determined from TDP43 LC alone, *IC_50_* is the half maximal inhibitory concentration, *[H]* is the concentration of HSPB8, and *n* is the Hill coefficient, a measure of cooperativity.

### yADH Turbidity Assay

Yeast alcohol dehydrogenase (yADH, Sigma Aldrich) was dissolved in 50 mM sodium phosphate, pH 8.0, 100 mM sodium chloride and filtered with a 0.22 µm syringe filter. The concentration was determined by UV-Vis absorbance at 280 nm (E^1%^ = 14.6). HSPB8, in 25 mM Tris pH 8, 5 mM DTT and yADH were combined and incubated for 10 minutes at RT with 1 mM DTT, then diluted with 50 mM sodium phosphate, pH 8.0, 100 mM sodium chloride to achieve final concentrations of 0.2 mg/mL (9 μM monomeric) HSPB8, 0.125 mg/mL (90 μM tetrameric) yADH, and 20 mM DTT. The assay was performed using a 96 well plate sealed with Thermo Scientific Nunc PVC-coated polypropylene adhesive seals. A BioTEK Synergy H1 plate reader set to 42 °C with 360 nm absorbance readings taken every 30 s for 1 h total. Each condition was performed in triplicate with 90 µL in each well.

### SEC Assay

The concentration of HPSB8 was determined from the 280 nm absorbance on a ThermoScientific Nanodrop 2000c with a 1 mm pathlength and a calculated extinction coefficient (Protparam, Expasy). Samples were then buffer exchanged and concentrated as needed in an Amicon Ultra 0.5 mL 3K MWCO centrifugal filter with centrifugation at 14,000 g in 5–10 min intervals. The solution was mixed with a pipette between centrifugation cycles. The buffer for the protein and column was 150 mM NaCl, 20 mM Tris pH.8 or 150 mM sodium chloride, 20 mM HEPES pH. 7.5. The samples were centrifuged for 5 minutes at 21300 *g*, and 250 µl of each run over a Bio-Rad Enrich SEC 650 10 x 300 Column using a GE AKTA Pure 25 system.

### MALDI-TOF

TDP43 LC diluted to 40 µM and buffer exchanged into 20 mM ammonium formate, pH 3.0 using 3K MWCO Amicon Ultra-0.5 mL centrifugal filters with centrifugation at 14,000 g in 10 min intervals. Using the same procedure, HSPB8 was concentrated to 20 µM and buffer exchanged into 20 mM Ammonium Acetate, pH 7.2. For the combined TDP43 LC and HSPB8, they were mixed at a 1:2 molar ratio, in 20 mM Ammonium Acetate, pH 7.2 for 10–15 min before spotting.

Matrix solution was made by combining saturated 2,6-dihydroxyacetophenone in 96% ethanol and 18 mg/mL diammonium citrate in ultrapure water at a 3:1 ratio (v/v). MALDI-TOF samples were prepared by combining 2 µL of protein, 2 µL of 2% trifluoroacetic acid in ultra-pure water, and 2 µL of matrix solution. Finally, 1 µL of each sample was spotted on a 384-target ground steel plate and dried under vacuum for 15 min. All MALDI-TOF measurements were performed on a Bruker UltraFlextreme equipped with a 2 kHz SmartBeam laser. The instrument was calibrated using Protein Standard II (Bruker). The ion source 1 and 2 voltages were set to 20 kV and 18.25 kV, respectively. The lens detector voltage was set to 2.832 kV using a mass range of 10 to 50 kDa. The matrix suppression was done through deflection using a cut-off of 5 kDa. The spots were ionized by 250 shots in raster mode. Raw data was baseline corrected using flexAnalysis software (Bruker) then normalized and smoothed (Savitzky–Golay filter) in Python.

### Transmission Electron Microscopy

Samples were imaged using ultrathin carbon films (<3 nm) on 400 mesh lacey carbon copper grids (Ted Pella). The grids were prepared by adding 5 μL of sample and incubating for 2 min, then washing twice with 10 μL of ultrapure water for ∼10 s, and then stained with 5 μL of a 2% (w/v) uranyl acetate solution for 15 s. Between each step, bulk liquid was wicked away using a laboratory tissue prior to adding the next solution. Grids were imaged on a FEI Talos 120C transmission electron microscope operating at 80 keV.

### Fluorescent Labeling

Labeling reactions were conducted in 20 mM HEPES, pH 7.5, and 5 mM DTT by buffer exchanging TDP43 LC or HSPB8 using a centrifugal filter. Reactions used 20-fold molar excess of dye (Alexa Fluor 647 for TDP43 LC or NHS-Fluorescein for HSPB8, ThermoFisher Scientific) and 7.5–10 µM protein. For TDP43 LC, the labeling buffer also contained 6 M urea. Reactions were run in the dark at 4 °C under quiescent conditions for 18 h.

Reactions were then quenched by addition of 1 M Tris-HCl, pH 7.5 to 20 mM. Alexa Fluor 647-labeled TDP43 LC was diluted with 20 mM MES, pH 6.0, and 3 M Gu-HCl to 1 µM. Fluorescein-labeled HSPB8 was diluted with reaction buffer with 50% glycerol to 1–4 µM HSPB8 and 25% (v/v) glycerol. The HSPB8 alone FP samples were dialyzed back into reaction buffer in the dark prior to glycerol addition and freezing. Both proteins were aliquoted and stored at either –20 °C or –80 °C. Small quantities of the proteins were run on a non-reducing SDS-PAGE gel and imaged on a Bio-Rad ChemiDoc MP gel imager with presets for 647 and 488 nm to confirm labeling.

Before use in fluorescence experiments, fluorescently labeled proteins were thawed at 4 °C and desalted using 7k MWCO, 2 mL Zeba™ Spin Desalting Columns (ThermoFisher Scientific) into 6 M Urea, 10 mM PIPPS, pH 4.0 for TDP43 LC and 5 mM DTT, 20 mM Tris-HCl, pH 7.5 for HSPB8. The HSPB8 alone FP samples were dialyzed into the experimental buffer. This was done following the manufacturer’s recommendations. For FP experiments, labeled HSPB8 was used neat, for FRAP experiments the labeled proteins were pooled with stock unlabeled protein at a ratio of 1:300 for TDP43 LC and 1:100 for HSPB8 (labeled:unlabeled) and concentrated to 600 µM and 60 µM, following the procedure outlined in Thioflavin T Aggregation Assays.

### Fluorescence Polarization

His-cleaved, fluorescein labeled HSPB8 at 5 μM, TDP43 LC at 600 μM, 1 M Tris-HCl, pH 7.5, and ultrapure water were combined to achieve 300 µL of 5 nM HSPB8, 20 mM Tris-HCl, pH 7.5, 150 mM sodium chloride, 2 mM DTT, and various TDP43 LC or yADH concentrations. For triplicate measurements, 90 µL was aliquoted to each well of a black-bottom 96 well plate. Plates were pre-heated to 37 °C and unused outside wells were pre-filled with water following the same procedure described in Thioflavin T Aggregation Assays.

FP assays were performed on a 96 well black-bottomed plate sealed with Thermo Scientific Nunc PVC-coated polypropylene adhesive seals using a BioTEK Synergy H1 plate reader with parallel and perpendicular filters with excitations at 485 nm and emissions at 528 nm for both. Both sets of filters had 20 nm bandwidths and 510 nm dichroic mirrors. Samples were read from a height of 6 mm every 5 min, for 18 h, at 37 °C, with linear shaking for 30 s at the instrument’s maximum speed before each read.

For yADH, lyophilized protein was dissolved in 20 mM Tris HCl, pH 7.5 and mixed with Fluorescein labeled HSPB8 as described above. The samples were incubated at 37 °C for 15 min before the first read and 20.5 h for the second read.

### Fluorescence Polarization Data Analysis

The FP response at a given time, *t*, of labeled HSPB8 interacting with varying amounts of TDP43 LC that exhibit concentration independent aggregation rates (*k_app_*) is expected to take the general form:

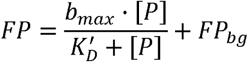

where *b_max_* is the apparent fluorescence polarization signal when HSPB8 saturates the available TDP43 LC binding sites (and is related to one or more identical binding sites on a TDP43 LC fibril), *[P]* is the concentration of TDP43 LC, *K′_D_* is the apparent equilibrium dissociation constant for HSPB8 binding to TDP43 LC, and *FP_bg_*is the apparent background fluorescence polarization signal. *K′_D_* is used rather than the dissociation constant (*K_D_*) to account for transient HSPB8-TDP43 LC interactions, and both *b_max_*and *FP_bg_* have time dependence.

FP values as a function of TDP43 LC concentration *at each time point* were fit to the above model using non-linear least squares analysis in Python. Standard deviations for each parameter were calculated from the diagonal elements of the covariance matrix obtained during the fitting process. Data points were weighted by their uncertainty for curve fitting purposes, using the following relationship:

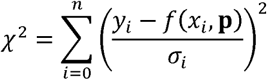

where χ*^2^* is the weighted sum of squared residuals, *y_i_* are the observed values, *f(x_i_,* **p**) are the predictions using vectorized model parameters, **p**, and σ*_i_* is the sample standard deviation of each data point. The best-fit parameters for *b_max_*, *K′_D_*, and *FP_bg_* were averaged for each time point to produce the thin lines and error bars in Fig. 2C.

Since these fitting parameters changed over time in the TDP43 LC-HSPB8 experiments, the measured values for these parameters were obtained by fitting the time-dependent averages to separate time-dependent functions. Final models were chosen based on the lowest Akaike Information Criterion and Bayesian Information Criterion.

For *b_max_,* a Monod model with quadratic decay was used:

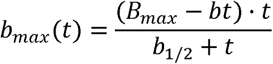

where, *t* is time, *B_max_* is the maximum fluorescence polarization signal for fully saturated HSPB8-TDP43 LC binding, *b* is the rate of non-recoverable loss of fluorescence polarization signal, and *b_1/2_*is the time required to achieve half of *B*_max_.

For *FP_bg_,* a sigmoidal model with linear decay was chosen:

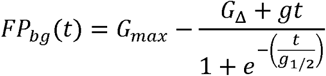

where, *G_max_* is the maximal background fluorescence polarization signal, *g* describes the rate of non-recoverable decrease in background FP over time, *g_1/2_* is the time to reach half minimal recoverable background fluorescence polarization signal. *G*_Δ_ is the change in background fluorescence between *G_max_*and the equilibrated background fluorescence signal (*G_bg_*) of the sigmoidal decrease in fluorescence polarization signal, described by:

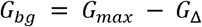

For *K′_D_*, the final model was a constant value and determined from the average of all data after background intensity reached equilibrium at 94 min. The thick lines in Fig. 2C were obtained from plotting the best fit *b_max_(t)* and *FP_bg_(t)* equations and the constant *K′_D_*.

The final FP binding curve plotted in Fig. 2D was obtained from:

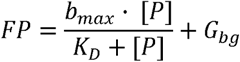

where *K_D_* the average *K′_D_* value after 94 min. To align with conventional reporting of FP binding curves, the background fluorescence, *G_bg_*, was excluded for the final plot.

### SDS-PAGE Staining and Imaging

Electrophoresis gels were fixed with 50 mL of water for 5 min then fixed and stained for 15 min with a solution containing 0.13% Coomassie Blue, 50% methanol, and 10% acetic acid. Gels were then washed with 100 mL of water and immediately destained twice for 1.5 h each with 25 mL of 12% methanol and 7% acetic acid. Gels were washed once more with 100 mL of water before imaging. Following Coomassie staining, silver staining was performed using a Pierce Silver Stain Kit (ThermoScientific). Washed gels were sensitized with gel sensitizer for 1 minute, washed with 50 mL of water, stained with silver stain for 30 minutes, washed with 50 mL of water, and finally the silver was reduced by developer for 2–3 min. The developer was quenched with 50 mL of 5% acetic acid for 10 min. Gels were washed with 100 mL of water before imaging using identical parameters. Images were taken on an ChemiDoc Imaging System (BioRad), using the white sample tray and Coomassie Blue or Silver Stain presets with auto-exposure.

### EDC Crosslinking

Pierce 1-ethyl-3-(3-dimethylaminopropyl)carbodiimide hydrochloride (EDC) No-Weigh^TM^ Format (ThermoFisher Scientific) and sodium;1-hydroxy-2,5-dioxopyrrolidine-3-sulfonate (sNHS, ThermoFisher Scientific) were dissolved in 20 mM MES to make 50 mM stock solutions. Stock solutions of proteins were prepared following the procedure outlined in Thioflavin T Aggregation Assays section. The final concentration for TDP43 LC was 600 µM and 60 µM for HSPB8.

TDP43 LC, EDC, and sNHS stock solutions were combined at a molar ratio of 1:50:100 to achieve final concentrations, 250 µM TDP43 LC, 12.5 mM EDC, and 25 mM sNHS and allowed to react for 15 min. Afterwards, the reaction was quenched with 1 M DTT stock solution to a final DTT concentration of 20 mM. HSPB8 was concentrated to 24 µM in 20 mM MES, pH 6.5, and combined with 1:50:100 EDC and sNHs and allowed to react for 15 min. Then the sNHS-activated HSPB8 was buffer exchanged into 20 mM HEPES, pH 8.0 and 5 mM DTT using 7k MWCO, 2 mL Zeba™ Spin Desalting Columns (ThermoFisher Scientific). The NHS-labeled TDP43 LC was then diluted with 22 µM sNHS-labeled HSPB8 in 20 mM HEPES, pH 8.0 and 5 mM DTT at a ratio of 1:12.5 (v/v) giving final concentrations of 20 µM for both TDP43 LC and HSPB8. Proteins were crosslinked at room temperature, under quiescent conditions for 30 min. The crosslinking reactions were quenched with 1 M Tris-HCl, pH 7.5 stock solution to a final concentration of 20 mM Tris-HCl. Crosslinked proteins were analyzed by SDS-PAGE.

### Amine-Reactive Crosslinking

Pierce bis(sulfosuccinimidyl) suberate (BS3) No-Weigh^TM^ Format (ThermoFisher Scientific) was dissolved in 20 mM HEPES pH 7.5 to make a 50 mM stock solution. Glutaraldehyde stock solutions at 2.6 M were diluted to 50 mM in 20 mM HEPES pH 7.5 immediately before use. TDP43 LC and HSPB8 were prepared following the procedure outlined in Thioflavin T Aggregation Assays to starting concentrations of 600 µM and 60 µM, respectively. Protein solutions were diluted to 20 µM with 20 mM HEPES, pH 7.5 at a molar ratio of 1:50 (protein to crosslinker) and allowed to react for 30 min. The reactions were then quenched with 1 M Tris-HCl, pH 7.5 to a final concentration of 20 mM Tris-HCl. Crosslinked proteins were analyzed by SDS-PAGE.

### Protein Digestion and Peptide Desalting

Quenched crosslinking reactions were buffer exchanged using 7k MWCO, 2 mL Zeba™ Spin Desalting Columns (ThermoFisher Scientific) into 1 M Gu-HCl, 20 mM Tris-HCl, pH 7.5. Reactions were reduced by the addition of 0.5 M TCEP to 5 mM and heating to 56 °C for 30 min. Acetylation was performed by adding 500 mM iodoacetamide (IAA) to 10 mM and incubating in the dark at RT for 1 h. Acetylation was then quenched for 15 min at RT by the addition of 1 M DTT to 15 mM.

Acetylated proteins were then buffer exchanged using 10K MWCO Amicon Ultra-15 centrifugal filters with centrifugation at 3000 x g in 20 min intervals into 1 M Urea, 20 mM Tris, pH 7. Proteins were digested for 1 h at 37 °C after the addition of 1 mg/mL Trypsin (proteomics grade, Sigma-Aldrich) in 50 mM acetic acid at a 1:100 mass ratio (trypsin:protein). A secondary digestion was performed for 18 h at room temperature by the addition of 0.5 mg/mL chymotrypsin (sequencing grade, Promega) in 1 mM HCl at a 1:100 mass ratio (chymotrypsin:protein). SDS-PAGE was used to confirm complete digestion of the crosslinked proteins. Digested proteins were desalted and concentrated using C18, 0.6 uL bed volume, 10 µL tip volume ZipTip® Pipette Tips (Millipore Sigma) following the manufacturer’s recommendations. Desalted peptides were then lyophilized and resuspended in 0.1% trifluoroacetic acid (TFA).

### Mass Spectrometry

Peptides were fractionated over 40 min at 0.500 µl·min^-1^ on a PepSep, 150 µm x 25 cm, 1.5 µm particle size, 100 Å pore, C18 analytical column (Bruker), heated to 49 °C, with 6–99% variable gradient (Supplemental Fig. S9). One µg of total peptides was injected and analyzed on an Orbitrap Exploris 480 using a MS1 scan range of 350-1500 (m/z), 1 microscan, normalized automatic gain control (AGC) target at 10^6^, radio frequency (RF) lens at 45%, in monoisotopic precursor selection (MIPS) mode. The minimum intensity was set to 5000 and charge states were filtered to include only charges 2–6. A dynamic exclusion filter was set to exclude after 1 time, for 30 s, using a mass tolerance of ± 10 ppm. Data dependent MS/MS was used to filter peptides for fragmentation, using 30 scans and a 1.6 m/z isolation window. Peptides were fragmented using a normalized higher-energy collisional dissociation (HCD) energy of 30%, an AGC target of 4·10^5^, 1 microscan, 40 ms ion accumulation time, RF lens set to 50%, with the first mass cutoff set to 170 (m/z). Data was set to centroid and MS1 and MS2 resolutions are 60000 and 15000 FWHM at 200 m/z, respectively.

### XL-MS Data Processing

Crosslinked peptides were identified using pLink2 (91). Search parameters were 20 ppm precursor mass tolerance, 50 ppm fragment mass tolerance, a minimum of 6 and a maximum of 60 amino acids per chain, and a peptide mass minimum of 600 and maximum of 6000 Da per chain. The crosslinker option was either BS3 or EDC-DE. A fixed modification was added for carbamidomethy (C; 57.02146) and variable modifications were added for deamidation (N, Q; 0.984016), oxidation (M; 15.994915), carbamyl (K, N-terminus; 43.005814). Trypsin and chymotrypsin were defined as a single custom enzyme with cleavage at the C-terminal side of FMLWYRK, with cleavage ignored when P1′ is Proline. Filter tolerance was set to ± 10 ppm using a 1% false discovery rate (FDR), separated for each subset of crosslinks (crosslink, looplink, monolink).

### Fluorescence Recovery After Photobleaching

Fluorescent protein was prepared as described in *Fluorescence Labeling* section. Liquid droplets consisting of TDP43 LC with 1:300 (labeled:unlabeled) fluorescent protein were formed by diluting 600 μM TDP43 LC in 6 M Urea, 10 mM PIPPS, pH 4.0 30-fold to 20 µM TDP43 LC with a final buffer composition of 10% (w/v) PEG-8000, 150 mM sodium chloride, 20 mM Tris-HCl, pH 7.5. Droplets containing both TDP43 LC and HSPB8 were prepared by diluting 60 μM HSPB8 in 20 mM Tris-HCl, pH 7.5, 5 mM DTT composed of 1:100 (labeled: unlabeled) fluorescent HSPB8 to 20 µM into the same buffer used for TDP43 LC. For each condition, 100 µL was prepared. Droplet formation was confirmed by bright-field microscopy using an Olympus BX51 light microscope with differential interference contrast (DIC) filters, a 40× objective lens, and a Diagnostics Instruments RT Slider camera with 6-megapixel sampling.

The samples were imaged by placing a pre-greased (silicon high-vacuum grease, DOW) PTFE O-ring, 006 Dash Number, ⅛″ ID, and ¼″ OD (Grainer) onto a 24 mm x 50 mm rectangular, #1.5H precision cover glass (ThorLabs). Then, 15 µL of the droplet solution was pipetted into the center of the O-ring and a half coverslip was placed on top. A diagram for sample preparation is in Supplemental Fig S5.

Recovery After Photobleaching (FRAP) experiments were performed using a Zeiss LSM 980 Aryscan confocal microscope with a Plan-Apochromat 63x/1.4 Oil DIC M27 objective using the Zen software package. Bleaching on the green and red fluorescence channels used 488-nm and 639-nm laser lines, respectively. Digital gain was set to 1 and offset to was set to 0. For 639 nm only measurements, the laser power was set to 0.8 %. When performing dual laser experiments, the 639 nm channel was set to 1.0% and the 488 nm channel was set to 0.15%. Pre-bleach fluorescence intensity was measured for 4 s, each spot was then bleached 5 times over the course of 0.23 s, and the post-bleach intensity was measured every 1 s for 1 min of recovery time. Raw FRAP data was normalized by setting the pre-bleach intensity to 1 and the post-bleach to 0, averaged (n = 4), then analyzed in Python by least-squares fitting to the following function:

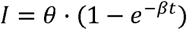

where *I* is the relative normalized intensity, *θ* is the fraction of mobile fluorophore which can be used to calculate the immobile fraction (α) by:

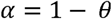

and β is related to the τ*_1/2_*, the time it takes to reach 50% of the mobile fraction intensity (θ), by:

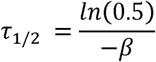

The supplemental bright-field and fluorescence images were captured on an Olympus IX81 inverted microscope equipped with a FV1000 Laser Scanning Confocal Microscope system. Images were taken with a 60x oil immersion objective (Olympus PlanApo N 60x/1.42 na Oil Objective). Fluorescent confocal imaging used an Argon laser with an excitation wavelength of 488 nm at 2.0%, HeNe laser with an excitation wavelength of 633 nm at 10.0%, and an excitation dichroic mirror set to 488/543/633 nm. Fluorescence emission was detected between 510 and 560 nm for the 488 nm line and between 650 and 750 nm for the 633 nm line. Bright field images were recorded on a second channel parallel to the one used to record the fluorescence images. Images were collected using a scan speed of 4.0 µs/pixel, a total area of 640 x 640 pixels.

## Author Contributions

K.M.J., K.C., D.C.F., K.M.O., C.C.S, and S.C., performed the experiments. D.T.M. acquired funding for the research and supervised the experiments. K.M.J. and D.T.M. analyzed data, wrote the manuscript, and conceptualized the experiments.

## Acknowledgments

We thank Dr. Steven L. McKnight and Dr. Masato Kato for providing the TDP43 LC plasmid, Dr. William Jewell for assistance with the Mass Spectrometry Core Facility at UC Davis (partially funded by National Institutes of Health Shared Instrumentation award S10OD025271 and S10OD018913), Dr. Bradley Shibata for assistance with the Bio-Electron Microscopy Core Facility at UC Davis, Dr. Gabriela Grigorean for helpful discussions and performing the LC-MS/MS experiments at the Proteomics Core Facility at UC Davis (National Institutes of Health Shared Instrumentation award S10OD026918-01A1), Dr. Thomas Wilkop for assistance with the Light Microscopy Core Facility at UC Davis (partially funded by the National Institutes of Health Shared Instrumentation award 1S10OD026702-01), Dr. Arpad Karsai for assistance with the Keck Spectral Imaging Core Facility at UC Davis, and Dr. Marie Heffern and Dr. Rebeca Fernandez for the use of UV-Vis instrumentation and providing reagents for XL-MS.

Research reported in this publication was supported by the National Institute of General Medical Sciences of the National Institutes of Health under award number R35GM142892 (D.T.M.). The entirety of the research in this publication was supported under this award except for core facility instrument acquisition and maintenance. The content is solely the responsibility of the authors and does not necessarily represent the official views of the National Institutes of Health.

