## Supplemental Data for "Small heat shock protein HSPB8 interacts with a pre-fibrillar TDP43 low complexity domain species to delay fibril formation"

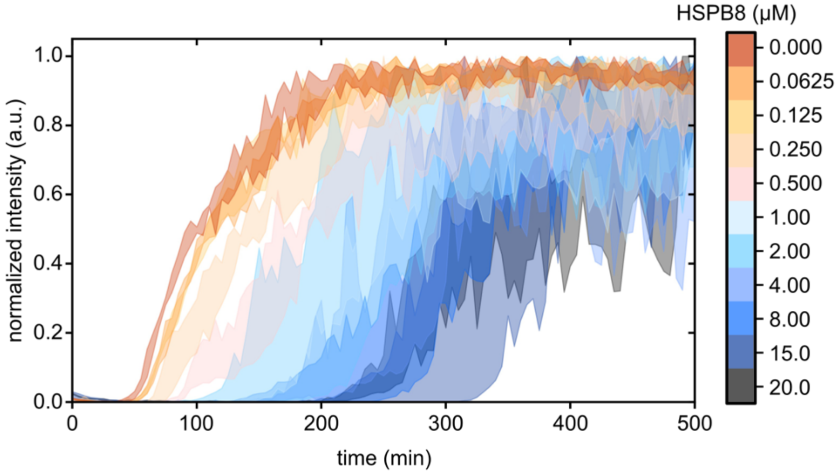


**Figure S1.** Normalized ThT fluorescence intensity traces for 20 µM TDP43 LC solutions with various concentrations of HSPB8. The filled regions represent the minimum and maximum of triplicate experiments.

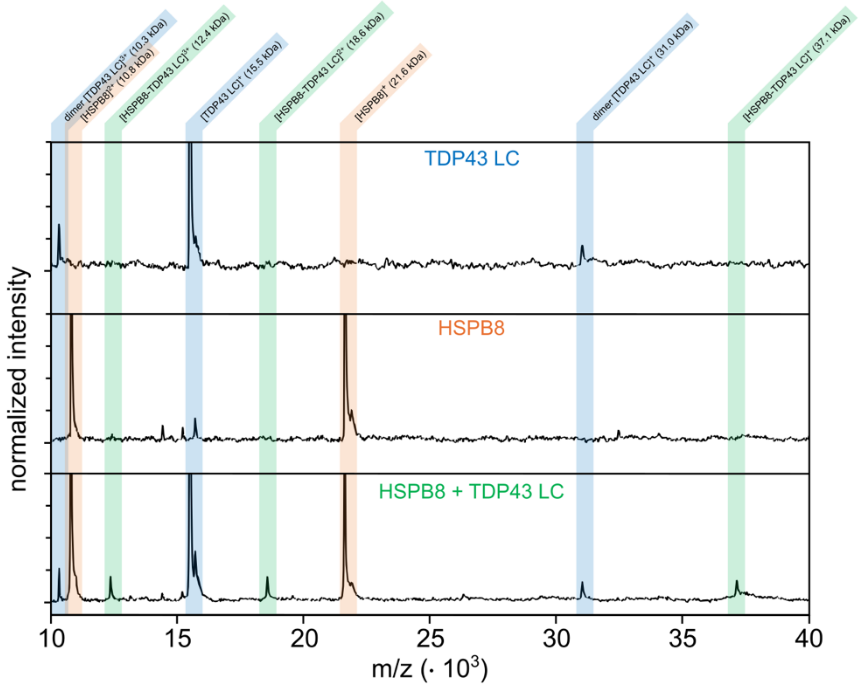


**Figure S2.** MALDI-TOF spectra of TDP43 LC alone (top), HSPB8 alone (middle), and HSPB8 with TDP43 LC at a 2:1 ratio (bottom). The mass spectrum of the mixture exhibits three new peaks (highlighted in the vertical green bars), corresponding to the $1^+$, $2^+$, and $3^+$ charge states of a 1:1 HSPB8-TDP43 LC complex. These peaks are not present in the isolated protein mass spectra, highlighted with blue (TDP43 LC) and orange (HSPB8) vertical bars.

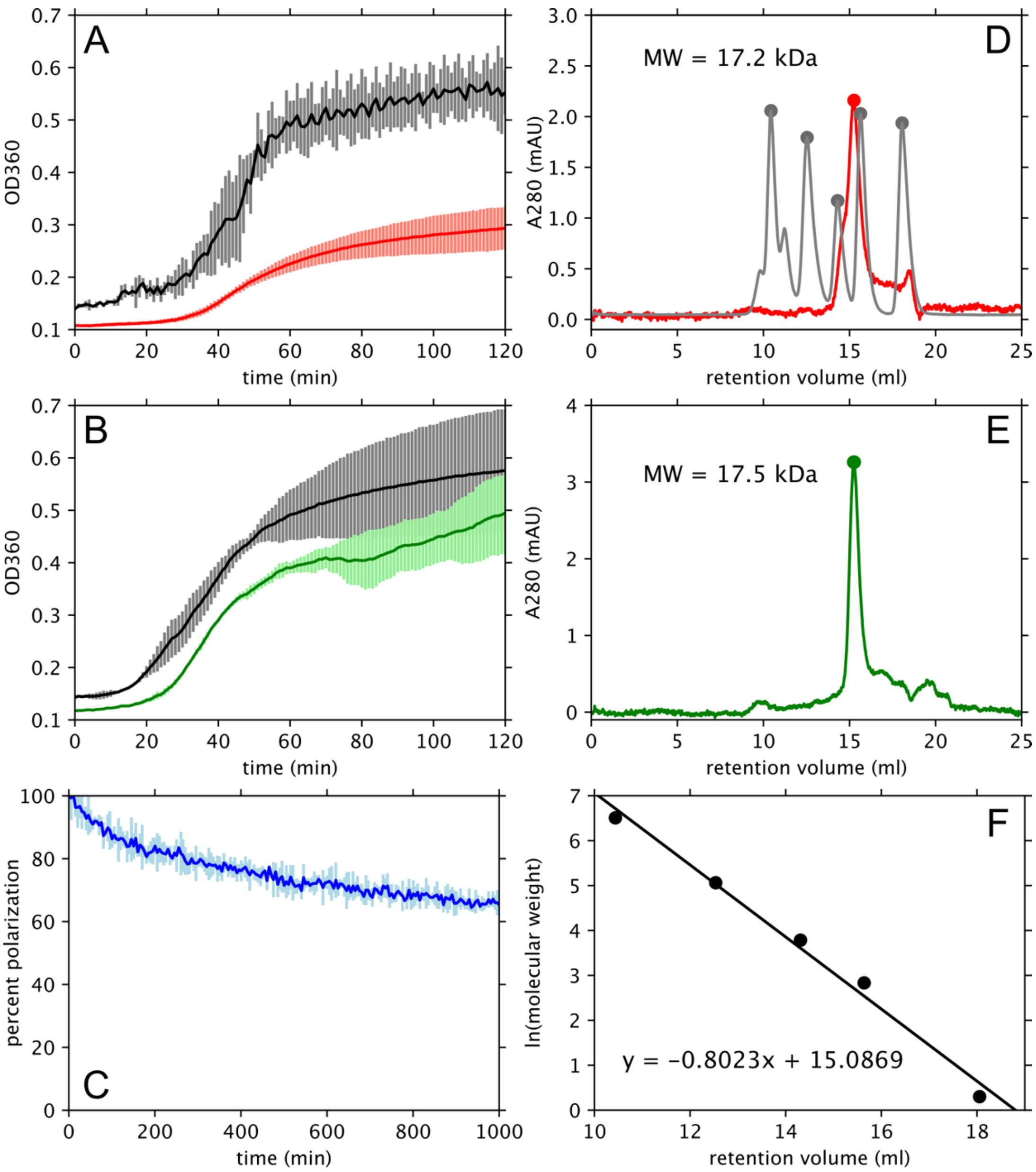


**Figure S3.** (A) Purified and His-tag cleaved HSPB8 delays DTT-induced aggregation of yeast alcohol dehydrogenase (yADH), measured by the increase in light scattering ($A_{360}$). The black line is the yADH alone and the red line is yADH with HSPB8. (B) The same assay as in panel (A) but with NHS-fluorescein labeled HSPB8 in green. (C) Fluorescence polarization (FP) signal, represented as a percentage of the initial value, of 5 μM NHS-fluorescein tagged HSPB8 alone at 37 °C, recorded under the same conditions as the FP assay of HSPB8 with TDP43-LC shown in Figure 3. For panels (A)–(C) the solid line is the average from three experiments on three different samples and the error bars represent ±1 standard deviation. (D) Size exclusion chromatography (SEC) of His-tag cleaved HSPB8 in buffer, red line, and standard proteins, gray lines. (E) SEC of NHS-fluorescein labeled HSPB8 in buffer. (F) Calibration plot for the SEC with best-fit line. The molecular weights stated in panels (D)–(E) are calculated using the best-fit line and the retention time of the HSPB8 peak center.

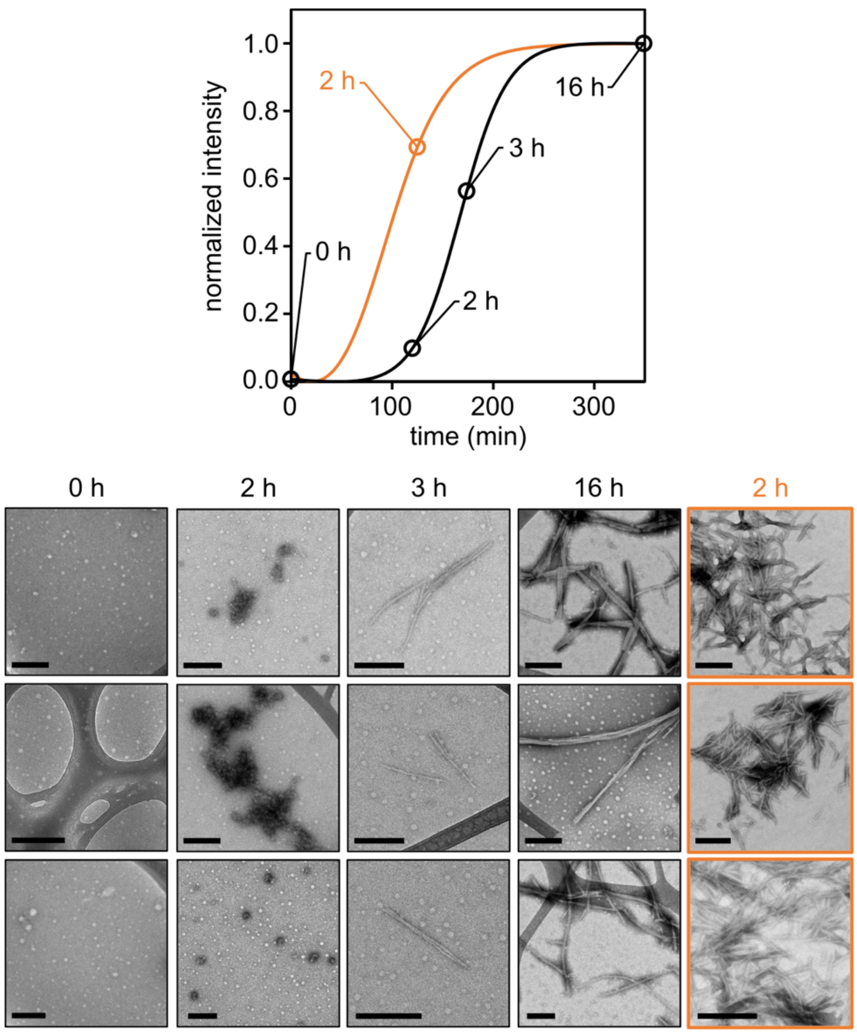


**Figure S4.** ThT curves of 20 μM TDP43 LC control (orange) or with 2 μM HSPB8 (black). Circled time points corresponded to the TEM images below. There are three images for each time point and all scale bars represent 250 nm. At the start (0 h) there are no signs of fibrils or aggregates. The small white spots are likely dust or other non-protein particles, as we have observed similar artifacts for TEM grids not containing HSPB8. At 2 h, amorphous aggregates begin to form in the presence of HSPB8, but short, clumped fibrils are already present in the negative control without HSPB8. At 3 h a few short, well dispersed fibrils are observed in the HSPB8 sample. Finally, after 16 h large, bundled fibrils have formed in the presence of HSPB8.

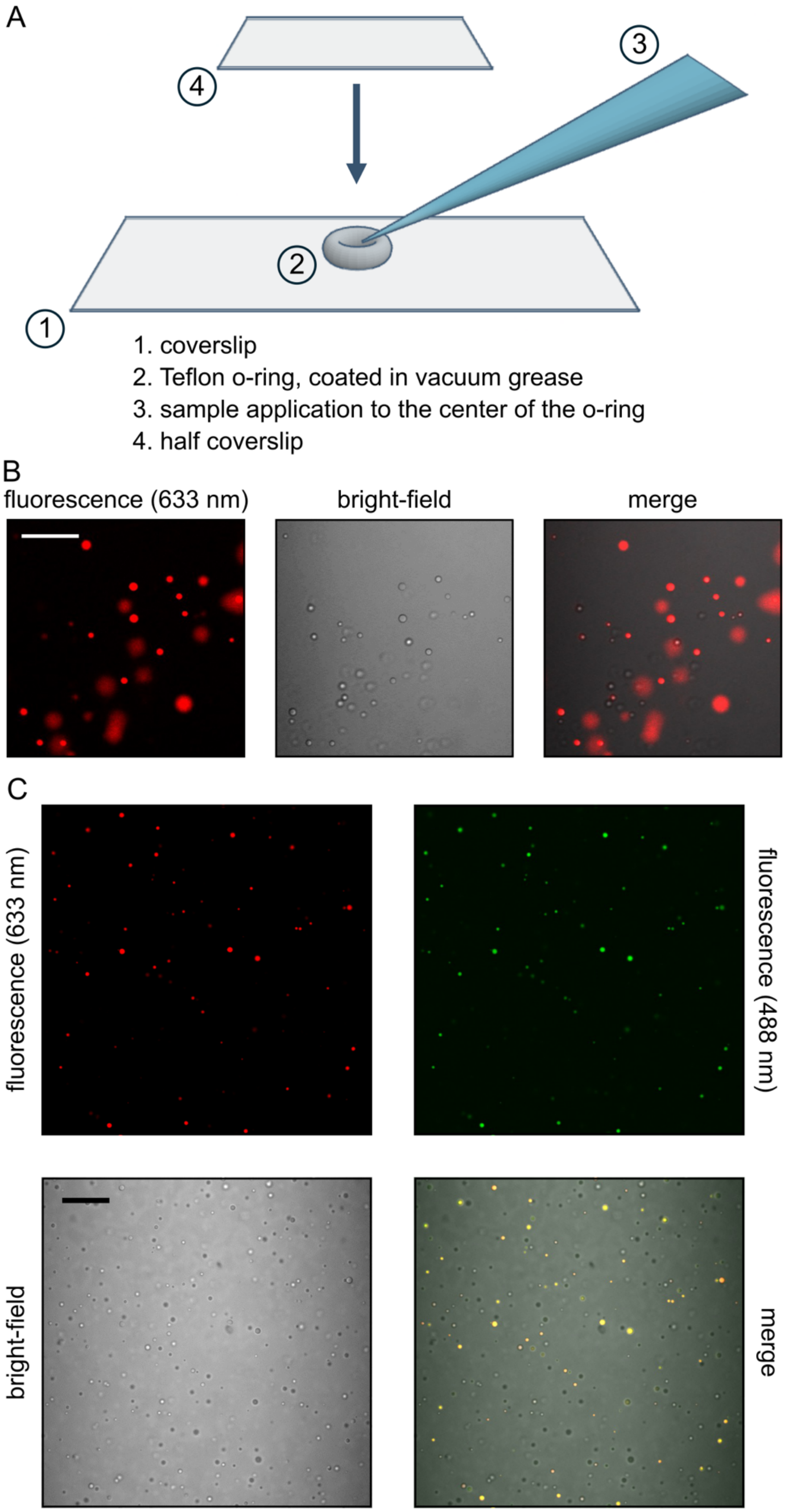
A
4
3
2
1
1. coverslip
2. Teflon o-ring, coated in vacuum grease
3. sample application to the center of the o-ring
4. half coverslip
B
fluorescence (633 nm)
bright-field
merge
C
fluorescence (633 nm)
fluorescence (488 nm)
bright-field
merge

**Figure S5.** (A) Representative diagram for FRAP sample preparation and imaging. A complete description is provided in the *Materials and Methods* section of the main text. (B) Confocal microscopy images showing TDP43 LC droplets. The diffuse areas of red are droplets that have settled to the surface of the microscope slide and immediately spread. Inset scale bar represents 30 μm. (C) confocal and bright field images of TDP43 LC droplets in the presence of HSPB8. Confocal microscopy images showing HSPB8 (green) co-localized in TDP43 LC droplets (red). The merged frame includes all three others. Inset scale bar represents 30 μm.

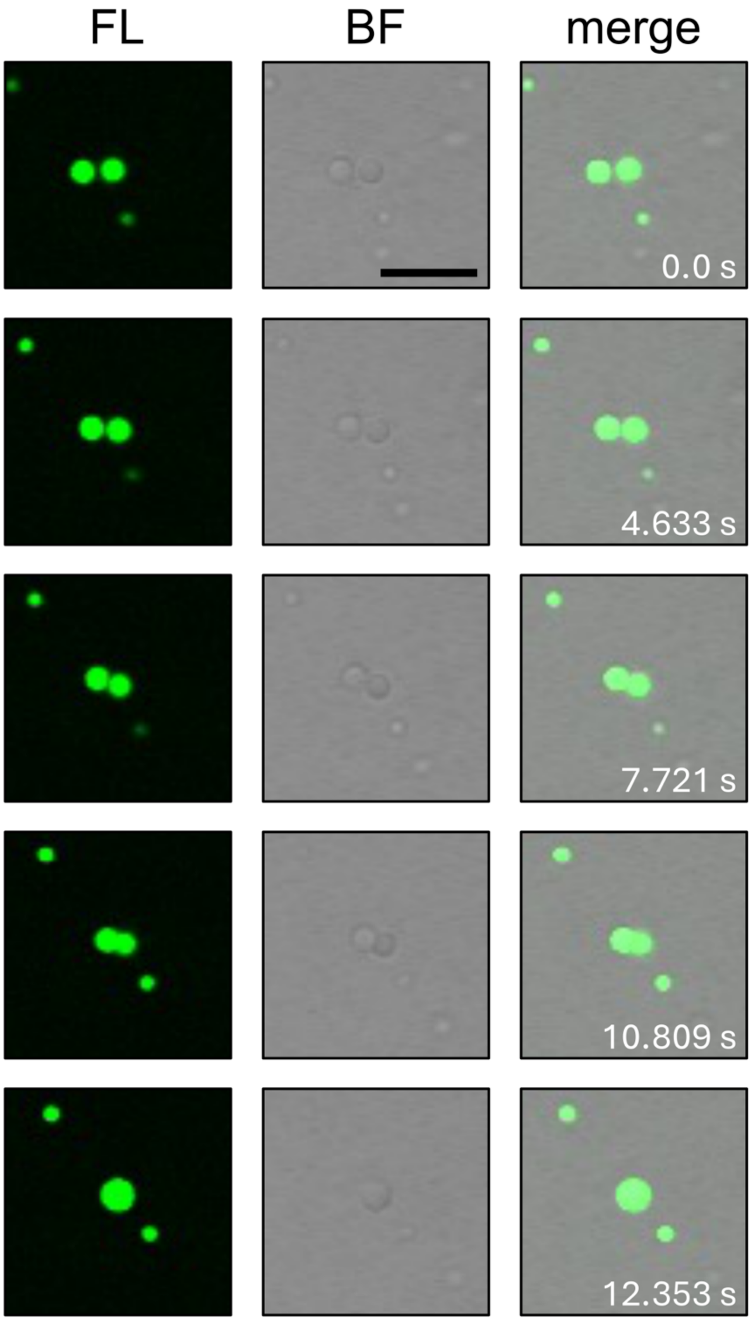


**Figure S6.** Confocal microscopy of TDP43 LC droplets in bulk solution in the presence of fluorescein-labeled HSPB8. FL is the fluorescent channel, BF is the brightfield channel, and Merge is the combined channel. The timepoint for each set of images is shown inset under the Merge column. Scale bar in the first BF image represents 10 μM and is the same for all images.

### TDP43 LC

```
-1         271        281        291        301        311        321
GAMD PKHNSNRQLE RSGRFGGNPG GFGNQGGFGN SRGGGAGLGN NQGSNMGGGM NFGAFSINPA

           331        341        351        361        371        381
     MMAAAQAALQ SSWGMMGMLA SQQNQSGPSG NNQNQGNMQR EPNQAFGSGN NSYSGSNSGA

           391        401        411 414
     AIGWGSASNA GSGSGFNGGF GSSMDSKSSG WGM
```

### HSPB8

```
 -1         10         20         30         40         50         60
 G   SADGQMPFSC HYPSRLRRDP FRDSPLSSRL LDDGFGMDPF PDDLTASWPD WALPRLSSAW

            70         80         90        100        110        120
     PGTLRSGMVP RGPTATARFG VPAEGRTPPP FPGEPWKVCV NVHSFKPEEL MVKTKDGYVE

           130        140        150        160        170        180
     VSGKHEEKQQ EGGIVSKNFT KKIQLPAEVD PVTVFASLSP EGLLIIEAPQ VPPYSTFGES

           190    196
     SFNNELPQDS QEVTCT
```

**Figure S7.** Sequences of the TDP43 LC and HspB8 constructs used in this paper. Underlined and bolded residues are overhangs leftover from the cleaved purification tag. The green highlighted residues are acidic residues that can be crosslinked by EDC. The yellow residues are basic residues that can be crosslinked to an acidic residue with EDC or another basic residue when crosslinked with BS3, DSS, or glutaraldehyde. Domains are highlighted to represent the transient $\alpha$-helix (orange), the aggregation core (red), and the a-crystallin domain (blue).

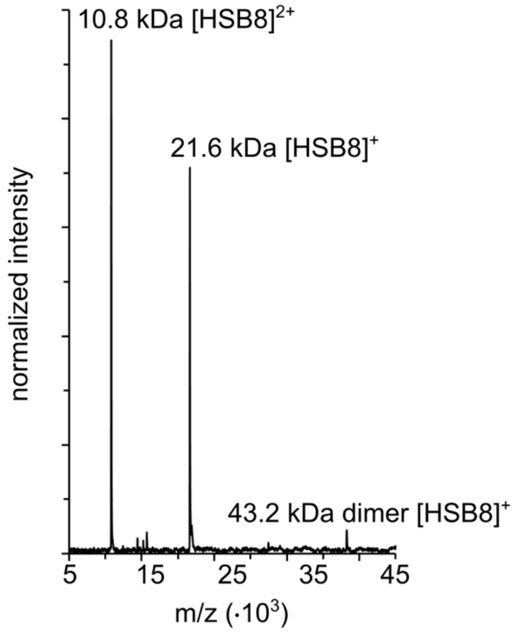


**Figure S8.** MALDI-TOF spectrum of HSPB8 in 10 mM ammonium carbonate, pH 7.2. A low intensity peak corresponding to the dimer is present.

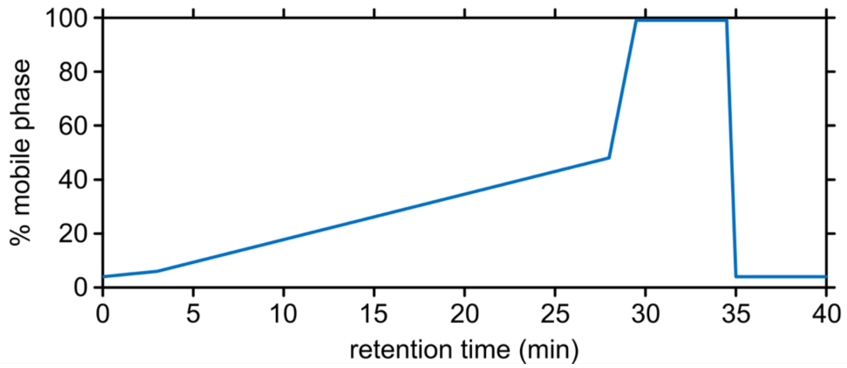


**Figure S9.** Peptide fractionation scheme. Elution from 3 to 39 min is directed to the mass spectrometer. Mobile phase A consists of 0.1% formic acid in water and mobile phase B is 80% acetonitrile, 0.1% formic acid in water.

For the following Tables S1-4, only the highest E-value scoring peptide is listed for each unique entry.

**Table S1.** List of EDC-crosslinked sites and corresponding peptides.

| Protein1 (Link1)-Protein2 (Link2) | Precursor Mass | Peptide 1 (Link1)-Peptide 2 (Link2) | Peptide Mass | E-value | Score |
|---|---|---|---|---|---|
| HSPB8 (124)-HSPB8 (131) | 2632.316405 | TKDGYVEVSGKHEEK(11)-QQEGGIVSK(3) | 2632.321177 | 8.47E-14 | 5.02E-02 |
| | 2893.427985 | TKDGYVEVSGKHEEK(11)-QQEGGIVSKNF(3) | 2893.432508 | 1.92E-11 | 1.91E-02 |
| | 2893.432163 | DGYVEVSGKHEEK(9)-QQEGGIVSKNFTK(3) | 2893.432508 | 3.80E-06 | 4.11E-02 |
| | 2664.289781 | DGYVEVSGKHEEK(9)-QQEGGIVSKNF(3) | 2664.289879 | 8.76E-09 | 5.60E-02 |
| | 3122.573781 | TKDGYVEVSGKHEEK(11)-QQEGGIVSKNFTK(3) | 3122.575137 | 8.50E-06 | 1.49E-01 |
| TDP-43 LC (362)-HSPB8 (116) | 2392.147057 | EPNQAF(1)-TKDGYVEVSGKHEEK(2) | 2392.141437 | 1.34E-05 | 1.53E-07 |
| | 1868.902406 | EPNQAF(1)-TKDGYVEVSGK(2) | 1868.9024 | 5.06E-05 | 7.39E-07 |
| | 3071.39917 | EPNQAFGSGNNSY(1)-TKDGYVEVSGKHEEK(2) | 3071.397576 | 6.02E-03 | 1.44E-03 |
| | 2153.067949 | QREPNQAF(3)-TKDGYVEVSGK(2) | 2153.062074 | 1.32E-01 | 5.26E-02 |
| HSPB8 (120)-HSPB8 (128) | 2632.317568 | TKDGYVEVSGK(7)-HEEKQQEGGIVSK(4) | 2632.321177 | 6.43E-13 | 7.35E-03 |
| | 2859.479267 | VKTKDGYVEVSGK(9)-HEEKQQEGGIVSK(4) | 2859.484541 | 8.49E-17 | 3.27E-03 |
| | 2893.426571 | TKDGYVEVSGK(7)-HEEKQQEGGIVSKNF(4) | 2893.432508 | 7.81E-08 | 1.21E-01 |
| | 3120.592164 | VKTKDGYVEVSGK(9)-HEEKQQEGGIVSKNF(4) | 3120.595872 | 6.79E-07 | 1.28E-01 |
| TDP-43 LC (362)-HSPB8 (1) | 2274.93329 | EPNQAFGSGNNSY(1)-GSADGQMPF(1) | 2274.935554 | 5.79E-04 | 3.84E-01 |
| | 1595.67684 | EPNQAF(1)-GSADGQMPF(1) | 1595.679416 | 7.43E-06 | 4.61E-02 |
| | 2559.097871 | QREPNQAFGSGNNSY(3)-GSADGQMPF(1) | 2559.095228 | 6.98E-03 | 2.01E-01 |
| TDP-43 LC (362)-HSPB8 (142) | 2526.33611 | QREPNQAF(3)-KIQLPAEVDPVTVF(1) | 2526.334983 | 3.74E-05 | 1.27E-01 |
| | 2242.172664 | EPNQAF(1)-KIQLPAEVDPVTVF(1) | 2242.175309 | 4.29E-03 | 4.76E-01 |
| | 2921.442132 | EPNQAFGSGNNSY(1)-KIQLPAEVDPVTVF(1) | 2921.431447 | 1.07E-03 | 3.21E-01 |
| TDP-43 LC (406)-HSPB8 (–1) | 2956.210672 | GSASNAGSGSGFNGGFGSSMDSK(21)-GSADGQMPF(1) | 2956.21071 | 1.47E-02 | 1.98E-02 |
| TDP-43 LC (362)-HSPB8 (137) | 2416.150641 | EPNQAF(1)-HEEKQQEGGIVSKNF(13) | 2416.15267 | 2.15E-03 | 7.12E-05 |
| TDP-43 LC (406)-HSPB8 (124) | 2815.363493 | DSKSSGW(1)-VEVSGKHEEKQQEGGIVSK(6) | 2815.38556 | 1.85E-03 | 5.68E-03 |
| TDP-43 LC (406)-HSPB8 (115) | 2250.031339 | NGGFGSSMDSK(9)-TKDGYVEVSGK(2) | 2250.034201 | 1.88E-03 | 5.81E-03 |
| TDP-43 LC (261)-HSPB8 (127) | 2437.176979 | DPKHNSNR(1)-DGYVEVSGKHEEK(12) | 2437.185363 | 4.05E-03 | 3.88E-01 |
| TDP-43 LC (–1)-TDP-43 LC (362) | 1912.87389 | GAMDPKHNSNR(1)-EPNQAF(1) | 1912.871783 | 5.32E-07 | 2.05E-02 |
| TDP-43 LC (362)-TDP-43 LC (150) | 2189.914549 | EPNQAF(1)-NGGFGSSMDSKSSGW(11) | 2189.919179 | 7.67E-03 | 4.49E-03 |
| | 2474.077814 | QREPNQAF(3)-NGGFGSSMDSKSSGW(11) | 2474.078853 | 1.47E-05 | 1.45E-02 |
| | 2098.923031 | QREPNQAF(3)-GSSMDSKSSGW(7) | 2098.924599 | 2.80E-07 | 8.66E-02 |
| | 1452.639098 | EPNQAF(1)-DSKSSGW(3) | 1452.638937 | 8.27E-05 | 3.50E-01 |
| HSPB8 (115)-HSPB8 (120) | 1933.000474 | VKTKDGY(4)-VEVSGKHEEK(2) | 1933.002442 | 4.54E-04 | 9.25E-02 |
| | 064.03882 | MVKTKDGY(5)-VEVSGKHEEK(2) | 2064.042921 | 5.17E-05 | 9.91E-02 |
| | 2990.527114 | MVKTKDGY(5)-VEVSGKHEEKQQEGGIVSK(2) | 2990.52502 | 3.00E-04 | 2.64E-01 |
| HSPB8 (127)-HSPB8 (137) | 2893.433942 | TKDGYVEVSGKHEEK(14)-QQEGGIVSKNF(9) | 2893.432508 | 2.35E-06 | 5.25E-03 |

| | | | | | |
|---|---|---|---|---|---|
| | 2664.289428 | DGYVEVSGKHEEK(12)-QQEGGIVSKNF(9) | 2664.289879 | 4.03E-08 | 5.98E-02 |
| TDP-43 LC (406)-HSPB8 (141)* | 2624.295477 | NGGFGSSMDSK(9)-KIQLPAEVDPVTVF(1) | 2624.291126 | 3.78E-07 | 8.15E-02 |
| TDP-43 LC (406)-HSPB8 (141) | 2477.297548 | GSSMDSK(5)-TKKIQLPAEVDPVTVF(2) | 2477.295486 | 6.97E-05 | 1.77E-01 |
| TDP-43 LC (362)-HSPB8 (124) | 3373.640389 | QREPNQAF(3)-DGYVEVSGKHEEKQQEGGIVSK(9) | 3373.640581 | 8.81E-02 | 2.19E-01 |
| TDP-43 LC (408)-HSPB8 (120) | 2815.358621 | DSKSSGW(3)-VEVSGKHEEKQQEGGIVSK(2) | 2815.38556 | 2.99E-02 | 3.87E-01 |
| TDP-43 LC (406)-HSPB8 (–1) | 2956.206322 | GSASNAGSGSGFNGGFGSSMDSK(21)-GSADGQMPF(1) | 2956.21071 | 5.85E-02 | 4.30E-01 |

* Only the deamidated peptide was found.

**Table S2.** List of EDC-crosslinked loops and corresponding peptides.

| Protein (Link1) (Link2) | Precursor Mass | Peptide Sequence (Link1) (Link2) | Peptide Mass | E-value | Score |
|---|---|---|---|---|---|
| HSPB8 (127)(128) | 2385.162635 | DGYVEVSGKHEEKQQEGGIVSK(12)(13) | 2385.167983 | 1.94E-13 | 8.11E-04 |
| HSPB8 (115)(116) | 1687.825188 | TKDGYVEVSGKHEEK(2)(3) | 1687.828513 | 7.22E-17 | 7.32E-02 |
| | 1164.588543 | TKDGYVEVSGK(2)(3) | 1164.589476 | 5.31E-06 | 1.70E-05 |
| HSPB8 (116)(124) | 2385.168284 | DGYVEVSGKHEEKQQEGGIVSK(1)(9) | 2385.167983 | 5.76E-14 | 1.42E-06 |
| | 1458.685653 | DGYVEVSGKHEEK(1)(9) | 1458.685884 | 3.49E-04 | 1.45E-03 |
| HSPB8 (–1)(3) | 1721.721415 | GSADGQMPFSCHYPSR(1)(4) | 1721.715811 | 8.40E-14 | 3.31E-03 |
| TDP-43 LC (406)(408) | 1485.604065 | NGGFGSSMDSKSSGW(9)(11) | 1485.606255 | 9.31E-11 | 2.07E-02 |
| | 1110.450301 | GSSMDSKSSGW(5)(7) | 1110.452001 | 5.76E-06 | 3.67E-03 |
| TDP-43 LC (261)(263) | 1208.557056 | GAMDPKHNSNR(4)(6) | 1208.558858 | 8.52E-09 | 1.55E-03 |
| | 949.459706 | DPKHNSNR(1)(3) | 949.459808 | 7.55E-11 | 1.84E-04 |
| HSPB8 (124)(126) | 1458.684421 | DGYVEVSGKHEEK(9)(11) | 1458.685884 | 2.15E-02 | 7.25E-03 |
| TDP-43 LC (258)(261) | 1208.557919 | GAMDPKHNSNR(1)(4) | 1208.558858 | 9.26E-09 | 6.80E-07 |
| HSPB8 (120)(124) | 2385.160108 | DGYVEVSGKHEEKQQEGGIVSK(5)(9) | 2385.167983 | 7.65E-18 | 1.03E-05 |
| HSPB8 (124)(127) | 2385.163986 | DGYVEVSGKHEEKQQEGGIVSK(9)(12) | 2385.167983 | 5.12E-05 | 1.31E-03 |
| HSPB8 (106)(108) | 955.529527 | KPEELMVK(1)(3) | 955.528069 | 1.11E-16 | 1.11E-01 |

**Table S3.** List of BS3-crosslinked sites and corresponding peptides.

| Protein1 (Link1)-Protein2 (Link2) | Precursor Mass | Peptide 1 (Link1)-Peptide 2 (Link2) | Peptide Mass | E-value | Score |
|---|---|---|---|---|---|
| HSPB8 (137)-HSPB8 (142) | 2899.544834 | QQEGGIVSKNF(9)-KIQLPAEVDPVTVF(1) | 2899.545017 | 3.43E-06 | 1.15E-10 |
| | 3422.790238 | HEEKQQEGGIVSKNF(13)-KIQLPAEVDPVTVF(1) | 3422.784054 | 1.90E-11 | 1.63E-07 |
| HSPB8 (–1)-HSPB8 (124) | 2523.133788 | GSADGQMPF(1)-DGYVEVSGKHEEK(9) | 2523.134299 | 3.28E-05 | 6.16E-10 |
| | 2752.287488 | GSADGQMPF(1)-TKDGYVEVSGKHEEK(11) | 2752.276929 | 1.25E-03 | 3.81E-08 |
| HSPB8 (115)-HSPB8 (142) | 2875.532461 | TKDGYVEVSGK(2)-KIQLPAEVDPVTVF(1) | 2875.533785 | 2.21E-08 | 1.04E-05 |
| | 3398.767717 | TKDGYVEVSGKHEEK(2)-KIQLPAEVDPVTVF(1) | 3398.772822 | 1.54E-09 | 3.59E-06 |
| HSPB8 (124)-HSPB8 (137) | 2820.367866 | DGYVEVSGKHEEK(9)-QQEGGIVSKNF(9) | 2820.368516 | 6.00E-07 | 6.11E-08 |
| HSPB8 (–1)-HSPB8 (137) | 2776.290991 | GSADGQMPF(1)-HEEKQQEGGIVSKNF(13) | 2776.288161 | 5.73E-07 | 2.87E-07 |
| TDP-43 LC (408)-HSPB8 (115) | 2823.256269 | NGGFGSSMDSKSSGW(11)-TKDGYVEVSGK(2) | 2823.277655 | 1.03E-06 | 5.29E-06 |
| | 3346.504042 | NGGFGSSMDSKSSGW(11)-TKDGYVEVSGKHEEK(2) | 3346.516692 | 5.41E-08 | 3.00E-05 |
| TDP-43 LC (408)-TDP-43 LC (406)* | 2769.13739 | NGGFGSSMDSKSSGW(11)-GSSMDSKSSGW(7) | 2769.14018 | 7.19E-03 | 1.52E-03 |
| | 3144.293414 | NGGFGSSMDSKSSGW(11)-NGGFGSSMDSKSSGW(11) | 3144.294433 | 4.75E-01 | 1.76E-01 |
| HSPB8 (137)-HSPB8 (137)* | 3073.518996 | HEEKQQEGGIVSKNF(13)-QQEGGIVSKNF(9) | 3073.522378 | 1.65E-03 | 2.30E-08 |
| HSPB8 (106)-HSPB8 (137) | 2317.210994 | KPEELMVK(1)-QQEGGIVSKNF(9) | 2317.210702 | 1.36E-06 | 4.38E-08 |
| HSPB8 (124)-HSPB8 (142) | 3169.630907 | DGYVEVSGKHEEK(9)-KIQLPAEVDPVTVF(1) | 3169.630193 | 1.87E-09 | 2.51E-05 |
| HSPB8 (–1)-HSPB8 (115) | 2229.034862 | GSADGQMPF(1)-TKDGYVEVSGK(2) | 2229.037892 | 1.46E-03 | 2.78E-04 |
| TDP-43 LC (263)-TDP-43 LC (408) | 2867.245025 | GAMDPKHNSNR(6)-NGGFGSSMDSKSSGW(11) | 2867.247037 | 1.24E-07 | 4.70E-02 |

*Homomeric interpeptide crosslinks provide evidence of self-association.

**Table S4.** List of BS3-crosslinked loops and corresponding peptides.

| Protein (Link1) (Link2) | Precursor Mass | Peptide Sequence (Link1) (Link2) | Peptide Mass | E-value | Score |
|---|---|---|---|---|---|
| HSPB8 (141)(142) | 1923.083064 | TKKIQLPAEVDPVTVF(2)(3) | 1923.083651 | 1.34E-18 | 1.34E-08 |
| | 2184.193552 | NFTKKIQLPAEVDPVTVF(4)(5) | 2184.194982 | 2.73E-20 | 2.55E-08 |
| HSPB8 (124)(128) | 2541.24783 | DGYVEVSGKHEEKQQEGGIVSK(9)(13) | 2541.24662 | 1.15E-31 | 3.74E-11 |
| HSPB8 (113)(115) | 1547.834607 | VKTKDGYVEVSGK(2)(4) | 1547.831478 | 2.39E-33 | 1.99E-07 |
| HSPB8 (137)(141) | 1701.915551 | QQEGGIVSKNFTKK(9)(13) | 1701.916931 | 2.32E-24 | 4.34E-06 |
| TDP-43 LC (258)(263) | 1364.636245 | GAMDPKHNSNR(1)(6) | 1364.637496 | 6.27E-21 | 5.37E-05 |
| HSPB8 (115)(124) | 1843.905472 | TKDGYVEVSGKHEEK(2)(11) | 1843.90715 | 9.23E-12 | 7.20E-05 |
| HSPB8 (106)(113) | 1340.748723 | KPEELMVKTK(1)(8) | 1340.749336 | 2.13E-05 | 7.21E-02 |
| HSPB8 (128)(137) | 1867.915809 | HEEKQQEGGIVSKNF(4)(13) | 1867.918383 | 1.01E-04 | 9.98E-02 |
